# Interstitial fluid transport dynamics predict glioblastoma invasion and progression

**DOI:** 10.1101/2025.03.12.642840

**Authors:** Cora M. Carman-Esparza, Caleb A. Stine, Naciye Atay, Kathryn M. Kingsmore, Maosen Wang, Ryan T. Woodall, Russell C. Rockne, Jessica J. Cunningham, Jennifer M. Munson

**Author notes:** **Co-corresponding Correspondence to:** Jennifer Munson, Ph.D., Professor, Fralin Biomedical Research Institute Virginia Tech, Room 1210, 4 Riverside Circle, Roanoke, VA 24016.

## Abstract

Glioblastoma is characterized by aggressive infiltration into surrounding brain tissue, hindering complete surgical resection and contributing to poor patient outcomes. Identifying tumor-specific invasion patterns is essential for advancing our understanding of glioblastoma progression and improving surgical and radiotherapeutic strategies. Here, we leverage in vivo dynamic contrast-enhanced magnetic resonance imaging (DCE-MRI) to noninvasively quantify interstitial fluid velocity, direction, and diffusion within and around glioblastomas. We introduce a novel vector-based pathline analysis to trace downstream accumulation of fluid flow originating from the tumor core, providing a spatially explicit perspective on local flow patterns. We find that localized fluid transport metrics predict glioblastoma invasion and progression, offering a new framework to non-invasively identify high-risk regions and guide targeted treatment approaches.

**One sentence summary:** Invasion and progression of glioblastoma can be predicted with interstitial fluid flow patterns via magnetic resonance imaging.

## Introduction

Glioblastoma (GBM) is the most common malignant primary brain tumor, with a median overall survival of just 15 months following diagnosis and a 5-year survival rate of only 5% [1]. Characterized by extensive infiltration into surrounding brain tissue, GBM’s diffuse invasion makes complete surgical resection difficult, leaving residual tumor cells that drive inevitable recurrence and poor patient outcomes [2]. Despite advances in clinical imaging, predicting the regions where glioblastoma cells invade beyond the tumor bulk remains a critical challenge. Identifying areas at high risk for invasion is essential for improving surgical and radiotherapeutic planning, enabling more aggressive treatment in invasive regions and greater preservation in lower-risk areas.

Interstitial fluid flow (IFF) is an ever-present phenomenon in the brain, crucial for maintaining normal physiological functions. During tumor growth, however, IFF is altered by elevated interstitial pressures generated within the growing tumor mass [3]. This heightened pressure drives abnormal fluid flow across the tumor’s margins. Studies in rodent GBM models have shown that surrogate tracers for IFF, such as Evans Blue dye, correspond to regions of cell invasion [4], and mechanisms such as CXCR4-CXCL12 signaling and autologous chemotaxis have been implicated in the IFF-driven invasion process [5, 6]. While these studies offer valuable insights, they rely on invasive techniques or tissue samples and do not provide a means to quantify these dynamics in a clinically relevant, in vivo setting. There is a pressing need for advanced, quantitative, and spatially explicit imaging metrics to link IFF to local cell invasion in patients.

Magnetic resonance imaging (MRI) is the standard imaging modality for GBM diagnosis and monitoring. Specifically, dynamic contrast-enhanced MRI (DCE-MRI) is widely used to delineate tumor boundaries based on areas of contrast enhancement, as the contrast agent leaks through the abnormal vasculature associated with GBM [7]. Current surgical resection margins are typically defined to include the contrast-enhancing tumor border and an additional 2 cm margin, an aggressive approach that disregards local spatial heterogeneities in tumor invasion. This static use of DCE-MRI ignores valuable dynamic information about how the contrast agent leaks into the parenchyma, which could provide deeper insights into the tumor microenvironment. Leveraging the Lymph4D algorithm, previously validated for analyzing fluid transport [8], we can noninvasively quantify interstitial fluid velocity, directionality, and diffusion from DCE-MRI scans, extracting biologically meaningful metrics directly from clinical imaging.

Given the known influence of fluid flow on glioblastoma invasion, spatially explicit patterns of interstitial fluid movement within and around tumors may correspond to regions of cell invasion. Identifying these patterns using in-vivo imaging could enable the noninvasive detection of invasive regions. In this study, we leverage quantitative analysis of DCE-MRI to examine local interstitial fluid velocity and diffusion. We introduce a novel vector-based pathline analysis to track fluid movement patterns originating from the tumor core into the surrounding parenchyma. To validate the predictive power of these metrics, we assess their correlation with histologically confirmed invading tumor cells and longitudinal tumor progression identified through MRI. The spatially explicit, image-based biomarkers of fluid transport dynamics presented here offer understanding of GBM invasion and progression and a significant advancement toward patient specific treatment planning and optimization of therapeutic strategies.

## Results

### Fluid flow metrics are spatially heterogeneous

We performed DCE-MRI on implanted murine gliomas as described in detail in the Materials and Methods. Using Lymph4D, IFF magnitude, diffusion, and direction were calculated for each pixel within the tumor boundary and the surrounding parenchyma with measurable contrast agent. A heat map and distribution of the magnitudes for GL261 Mouse 1 shows local variation through the tumor bulk and surrounding parenchyma and highlights intra-mouse variability (**Figure 1A-B**). There is also significant inter-mouse variability in both the GL261 and GSC mice (**Figure 1C-D**). This variability is also true of the diffusion coefficient (**Figure 1E-H**). Using the directionality of flow, the trajectory of flow originating within the tumor is followed until termination (**Figure 1I**). For each region in the surrounding parenchyma, the total number of tumor-originating pathlines passing through that region was summed, generating a “tumor-originating pathline density” map. This metric also exhibits significant heterogeneity of the distribution of tumor originating fluid flow within and between mice (**Figure 1J-L**).

**Figure 1:**
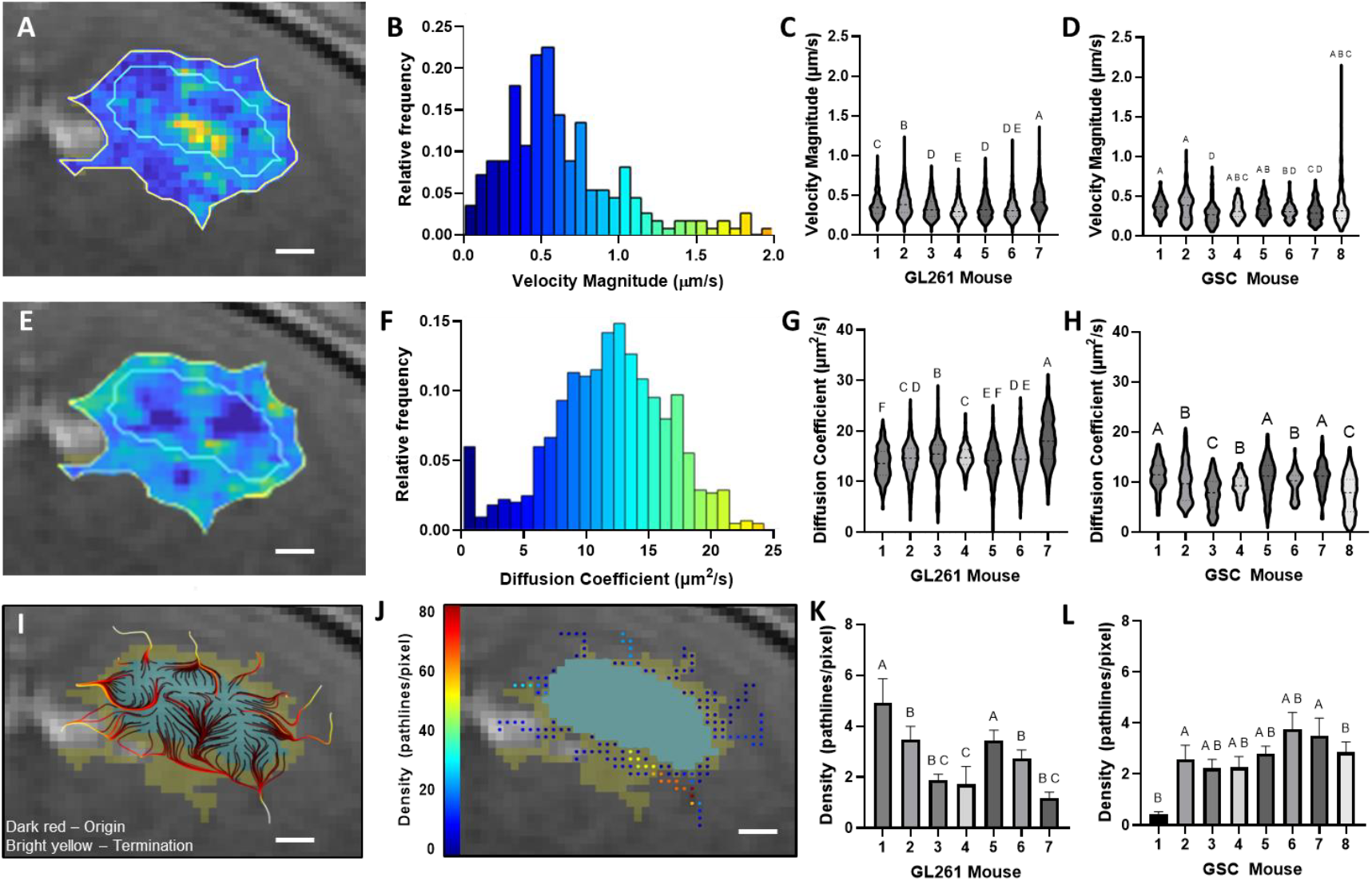
Intra- and inter-mouse variability of magnitude, diffusion, and tumor originating pathline density. A) Heat map of tumor and surrounding parenchyma IFF magnitude (um/s) with the tumor (blue boundary) and contrast-enhanced parenchyma (yellow boundary) (Scale bar =500μm). B) Distribution of magnitudes within GL261 Mouse 1. C) IFF magnitude of all pixels within the surrounding parenchyma for each GL261 mouse. Compact letter display shows columns are statistically indistinguishable if and only if they share at least one letter. D) IFF magnitude of all pixels within the surrounding parenchyma for each GSC mouse. E) Heat map of tumor and surrounding parenchyma IFF diffusion coefficient (um^2^/s) (Scale bar =500μm). F) Distribution of diffusion coefficients within GL261 Mouse 1. G) IFF diffusion coefficient of all pixels within the surrounding parenchyma for each GL261 mouse. H) IFF diffusion coefficient of all pixels within the surrounding parenchyma for each GSC mouse. I) Tumor-originating pathlines (dark red is initial location moving to ending location as bright yellow) (Scale bar =500μm). J) Density heatmap of tumor-originating pathlines where blue and red represent a low and high tumor-originating pathline density, respectively (Scale bar =500μm). K) Tumor originating pathline density of all pixels within the surrounding parenchyma for each GL261 mouse or location. L) Tumor originating pathline density of all pixels within the surrounding parenchyma for each GSC mouse or location.

### Multimodal co-registration allows for MRI derived flow metrics and IHC cell center analysis

Both MRI imaging and IHC sectioning were performed in the coronal plane (**Figure 2AB**). For each mouse, one or two MRI slices with the largest tumor bulk were selected (**Figure 2C**). Features such as ventricles, white matter, and tumor shape were used to identify two to three corresponding histological sections for each MRI slice. Tumor boundaries were determined on both the MRI and IHC slices (Methods), and the locations of invading cells were identified (**Figure 2D**). MRI and histological slices were co-registered using a multimodal registration approach (**Figure 2E**, Methods), enabling the spatial coordinates of invading GL261 cells identified by IHC to be directly mapped onto the MRI images (**Figure 2F**). This allowed for the integration of MRI-derived flow metrics with the precise locations of individual invading cells for spatially detailed analysis.

**Figure 2:**
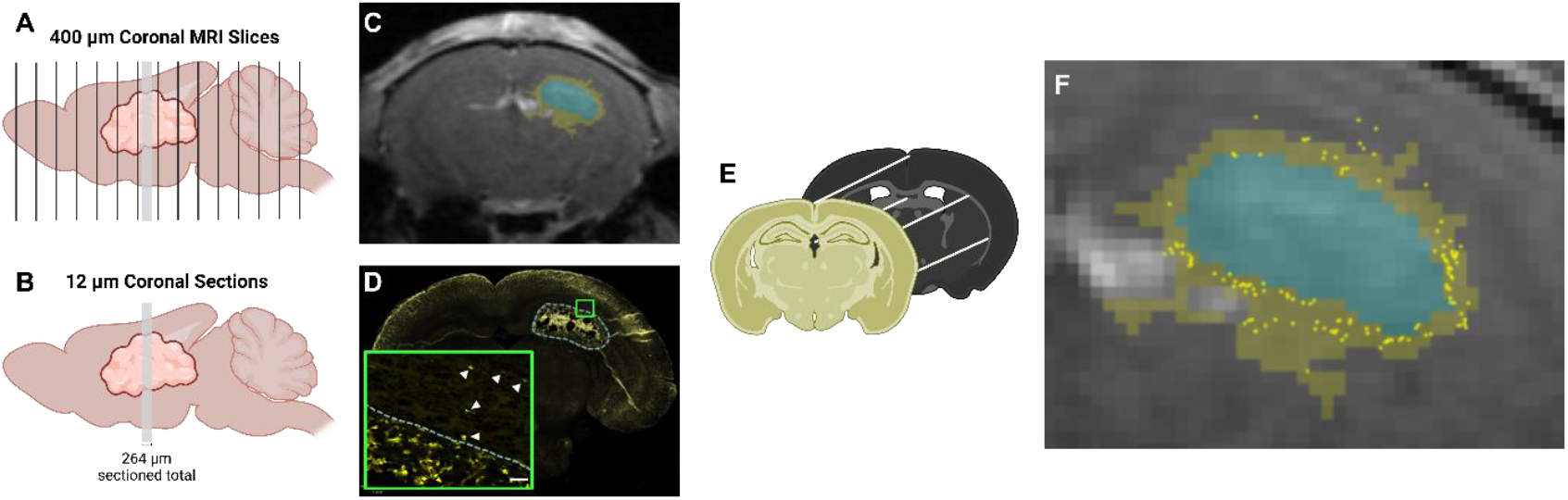
Multimodal co-registration of MRI and IHC cell centers. A) 400 μm coronal MRI slices through the entire murine brain. B) 12 μm IHC slices within the tumor bulk. C) MRI slice with mass tumor bulk, with the tumor (blue) and contrast-enhanced parenchymal (yellow) regions. D) Corresponding IHC slice with tumor boundary and inlay showing individual identified invading cells (white arrows). E) Multimodal registration using control point and geometric transformation. F) Combined MRI features and individual invading cell locations.

### Average tumor velocity magnitude is positively correlated with overall invasiveness

IFF velocity magnitudes were measured in the tumor and the surrounding contrast-enhanced parenchyma of implanted GL261 tumors (**Figure 3A**). The average velocity within the tumor is positively correlated with average velocity of the parenchyma (Spearman r = 0.7509, p-value = 0.0066). The average velocity magnitude in the tumor and surrounding parenchyma for each mouse location was compared to the total number of invading cells counted in the matching histological slices for that MRI location (**Figure 3B**). Velocity magnitudes in the tumor were significantly correlated with the number of invading tumor cells (Spearman r = 0.7509, p-value = 0.0066) (**Figure 3C**). The average velocity of the contrast-enhancing parenchyma and number of invading cells were not significantly correlated (Spearman r = 0.4807, p-value = n.s.) (**Supplemental Figure 5**).

**Figure 3:**
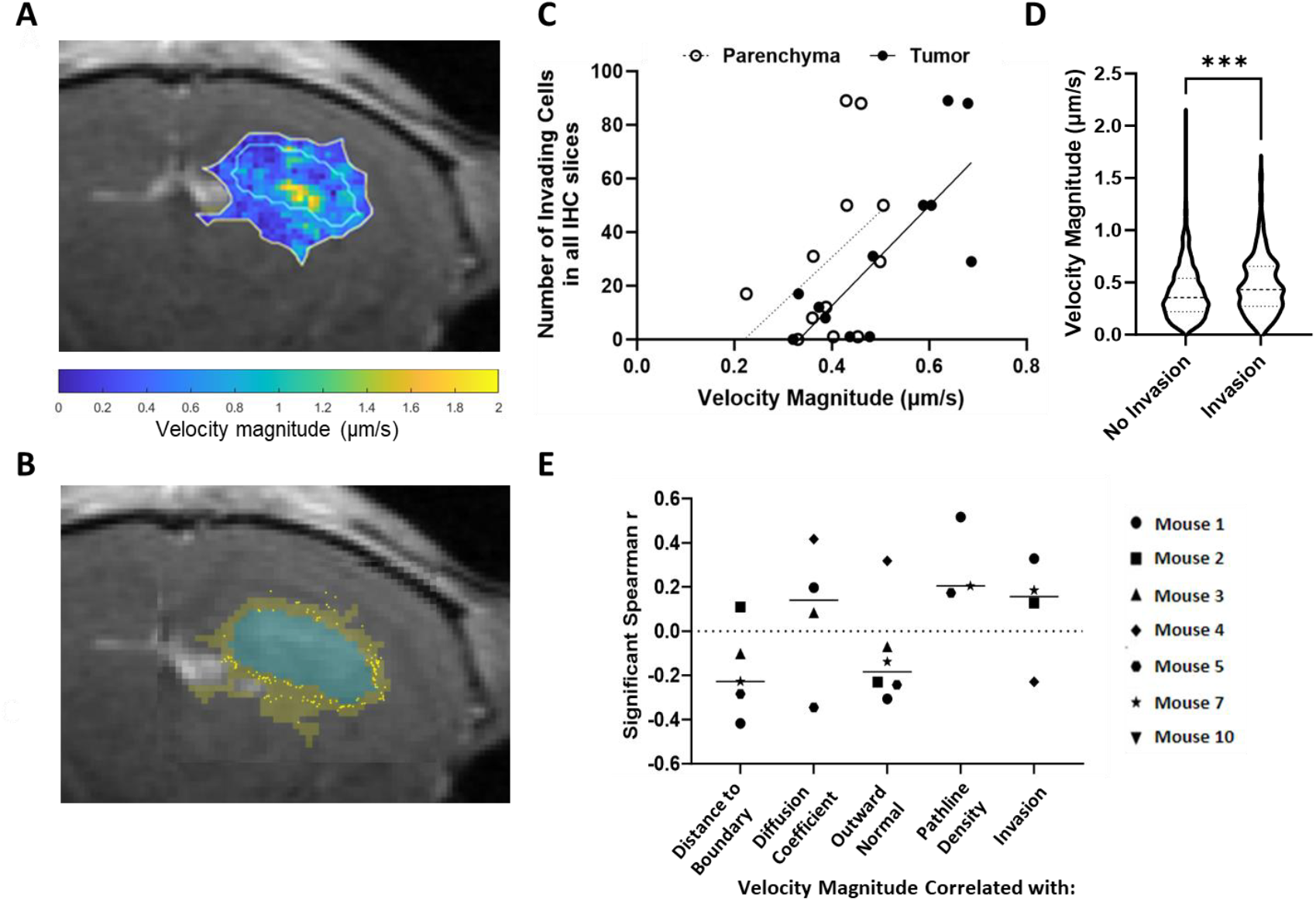
Increased velocity magnitude is associated with elevated invasion. A) Heatmap of velocity magnitude with tumor and contrast-enhancing parenchyma in light blue and yellow outlines, respectively (scale bar = 500 μm). B) Invasive cells (yellow points) overlaid on MRI (scale bar = 500μm). C) Correlation of total number of invading cells and average velocity magnitude within the tumor (Spearman r = 0.7509, p-value = 0.0066) and parenchyma (Spearman r = 0.4807, p-value = n.s.) for all MRI slices examined. D) Velocity magnitude in each region with and without at least one invading tumor cell (Mann-Whitney U test performed, p-value < 0.001; Cohen’s D = 0.28; no invasion (N=2402); invasion (N=208)). E) Multiparametric correlation analysis of local velocity magnitudes and each flow parameter. All correlations shown are significant as obtained from a Spearman correlation. Mice not shown for a specific parameter were not significant.

### Local velocity magnitude is elevated in invasive regions

The parenchymal regions that have at least one invading cell have a significantly faster velocity magnitude than regions with no invading cells; 0.47μm/sec ± 0.28 and 0.40μm/sec ± 0.25, respectively (p-value < 0.001; Cohen’s D = 0.28) (**Figure 3D**). Velocity magnitude is negatively correlated with distance to boundary, meaning that in four of the seven mice higher velocities were found closer to the tumor boundary.

Velocity magnitude is negatively correlated with the flow angle difference to outward normal in five of the seven mice, and positively correlated with tumor-originating pathline density, in three of the seven mice, showing that regions with elevated velocities were found where flow was directed away from the tumor boundary. Additionally, velocity magnitude is positively correlated with diffusion and invasive cells in multiple mice (**Figure 3E**).

### Local diffusion coefficient is lower in invasive regions

Diffusion coefficients were measured in the tumor and surrounding parenchyma of implanted GL261 tumors (Mouse 1 shown) (**Figure 4A**). The average diffusion coefficient in the tumor and the contrast-enhanced parenchyma were significantly correlated (**Supplemental Figure 5**). The average diffusion coefficient in the tumor and surrounding parenchyma for each mouse location was compared to the total number of invading cells counted in matching histological slices for that MRI location (**Figure 4B**), but these were not significantly correlated (**Figure 4C**). More importantly, when the spatially local diffusion coefficients were analyzed, as opposed to the overall bulk averages, regions that have at least one invading cell do have a significantly lower diffusion coefficient than regions with no invading cells (11.59±5.73 vs. 13.87±4.96 (p-value = <0.001; Cohen’s D = 0.43) (**Figure 4D**). This underscores the importance of considering the spatial heterogeneity of transport metrics and focusing on individual local regions rather than relying on overall averages. The diffusion coefficient is positively correlated with the distance to boundary for five of the seven mice, meaning that for each of these mice, the diffusion coefficient increases as the distance from the tumor increases (**Figure 4E**). There are only sparse significant correlations of diffusion with other IFF metrics across the mice, though five of the seven mice had significant negative correlation to invasion, which highlights that regions with invading cells have lower diffusion coefficients across the majority of the mice.

**Figure 4:**
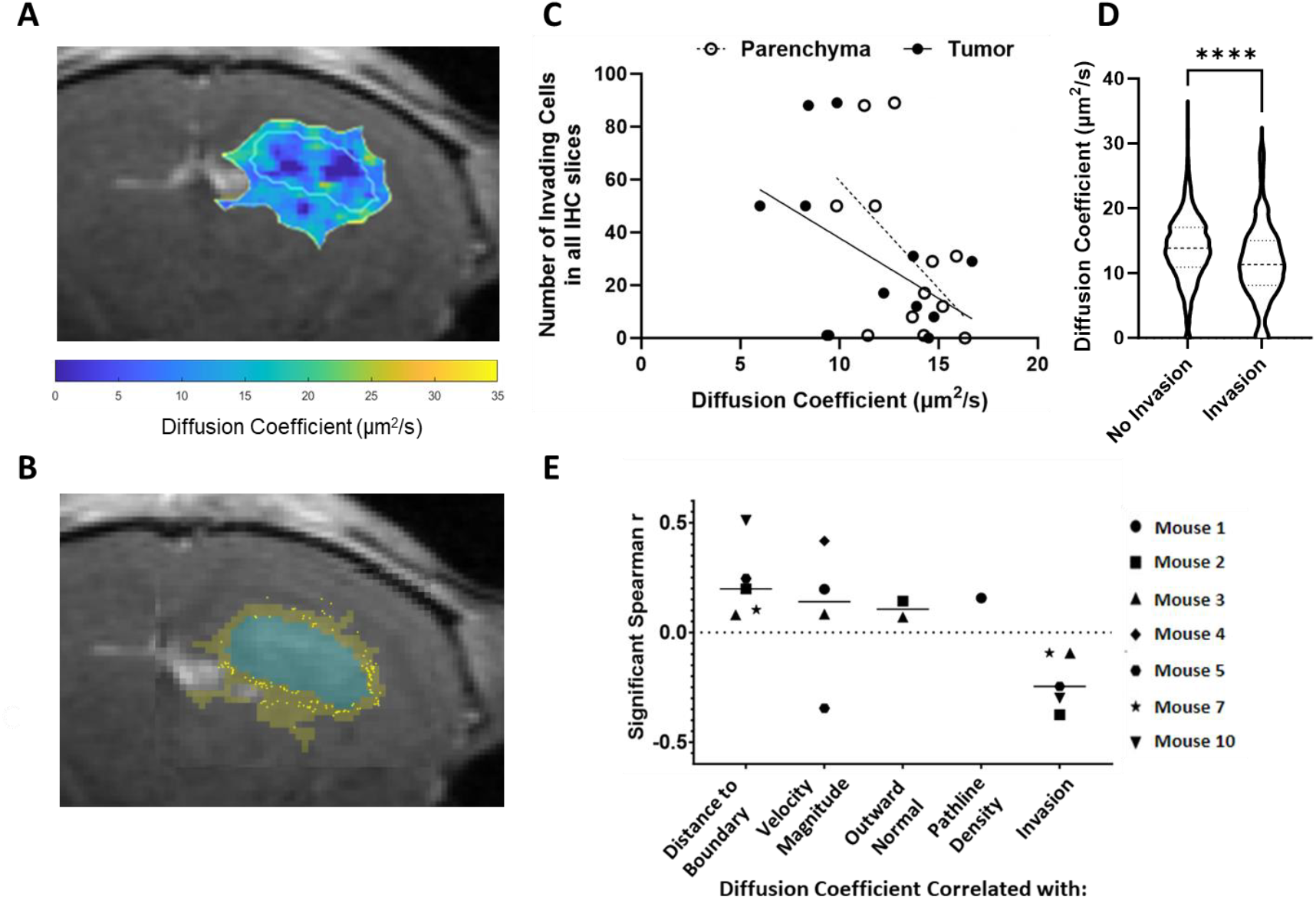
Decreased diffusion is associated with invasion. A) Representative heatmap of diffusion coefficient indicating the tumor and surrounding parenchyma in blue and yellow outlines, respectively (scale bar = 500μm). B) Invasive cells (yellow points) overlaid on MRI highlighting the tumor (blue) and contrast-enhanced parenchymal (yellow) regions (scale bar = 500μm). C) Average diffusion coefficient of the tumor and surrounding parenchyma is not significantly correlated with the number of invasive cells counted, though there is a negative trend (Tumor: Spearman r = -0.4351, p-value = n.s.; Parenchyma: Spearman r = -0.4737, p-value = n.s.). D) The diffusion coefficient in each local region with at least one invading tumor cell is significantly lower that regions without invasion (Mann-Whitney U test performed, p-value = < 0.001; Cohen’s D = 0.43; No Invasion (N=2402) Invasion (N=208)). E) Multiparametric correlation analysis of diffusion coefficient. All correlations shown are significant (p-value < 0.05) as obtained from a Spearman correlation. Mice not shown for a specific parameter were not significant.

### Tumor-originating fluid flow is significantly elevated in regions of invasion

The tumor-originating pathline density metric represents the relative amount of fluid originating from the tumor that traverses a given local region, with a higher pathline density indicating a greater volume of tumor-derived fluid passing through that parenchymal area. In Mouse 1, the tumor pathline density and histologically identified invading cells are shown. **(Figure 5A-C)**. Tumor-originating pathline density is significantly higher in local regions that have at least one invading cell (2.35±7.54 vs. 5.55±11.31 (p-value<0.001; Cohen’s D = 0.33) **(Figure 5D)**. This demonstrates that regions experiencing outward fluid flow from the tumor bulk into the parenchyma have significantly more invading cells, as quantified by this novel metric. Overall, we found that tumor originating pathlines intersected with 40-89% of all invading cells within each mouse with an average across all mice of 60.14% of invading cells (**Figure 5E**).

**Figure 5:**
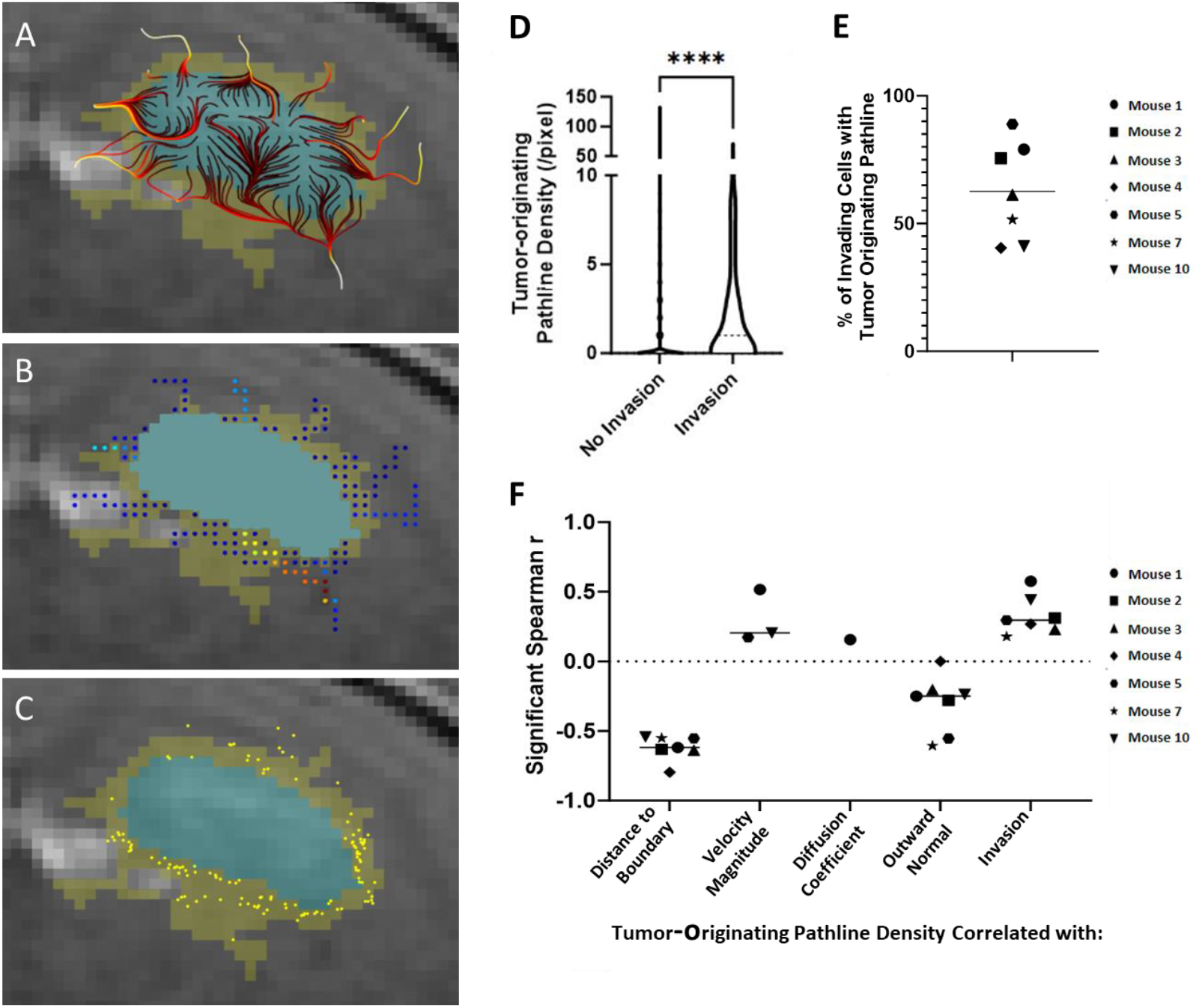
Tumor-originating pathline density is significantly elevated in regions of invasion. A) Tumor-originating pathlines overlaid on T1-weighted contrast-enhanced MRI with tumor (blue) and contrast-enhanced parenchyma (yellow). Dark red and yellow denote the original and ending position of tumor-originating pathlines, respectively (scale bar = 500 μm). B) Tumor-originating pathline density overlaid on T1-weighted contrast-enhanced MRI (dark blue and red represent a low and high number of tumor-originating pathlines, respectively) (scale bar = 500 μm). C) Invasive cells (dark yellow points) overlaid on a T1-weighted contrast-enhanced MRI (scale bar = 500 μm). D) Local tumor-originating pathline density with and without at least one invading tumor cell (Mann-Whitney U test performed, p-value < 0.001; Cohen’s D = 0.33; Invasion (N=2402) No Invasion (N=208)). E) Percentage within each mouse of invading cells with at least one tumor originating pathline within its region. F) Multiparametric correlation analysis of tumor-originating pathline density. All correlations shown are significant as obtained from a Spearman correlation.

In all seven mice, tumor-originating pathline density was negatively correlated with distance to the boundary, as expected, as all pathlines originate within the tumor, and the density of outward flow naturally diminishes with increasing distance from the tumor. In three mice there was a positive correlation with velocity magnitude meaning in these mice, regions with higher levels of tumor originating flow passing through them also experience higher velocity magnitudes. In all seven mice, outward flow is significantly correlated with invasion. Observing this pattern consistently across all mice in the study underscores the predictive power of this novel metric in identifying regions of invasion.

### Transport metrics significantly correlated with MRI-identified progressive disease

While identifying individual invading cells provides localized insights, radiographic progression reflects broader tumor growth dynamics. Here we analyze the underlying transport metrics associated not with individual cell invasion but with radiographic progression. For Mouse 1, the tumor boundary on Day 6 post injection (**Figure 6A**) and the tumor boundary on Day 8 post injection (**Figure 6B**) are shown. Creating a difference map between Day 8 and Day 6 highlights the pixels corresponding to areas of radiographic progression (purple highlight) with contrast-enhanced parenchyma (yellow) (**Figure 6C**). The tumor-originating pathline densities from the fluid flow measured on Day 6 are shown **(Figure 6D, Supplemental Dataset 2)**. We found that regions containing progression by Day 8 had a significantly faster velocity magnitude on Day 6 than regions that did not show progression (p-value = 0.0093; Cohen’s D = 0.15) (**Figure 6E**). Additionally, regions containing progression by Day 8 had a significantly lower diffusion coefficient on Day 6 than regions that did not show progression (p-value < 0.001; Cohen’s D = 0.26) **(Figure 6F, Supplemental Figure 3)**. Further, tumor-originating pathline density was significantly higher in regions that exhibited progression (p-value < 0.001; Cohen’s D = 0.31) **(Figure 6G)**.

**Figure 6:**
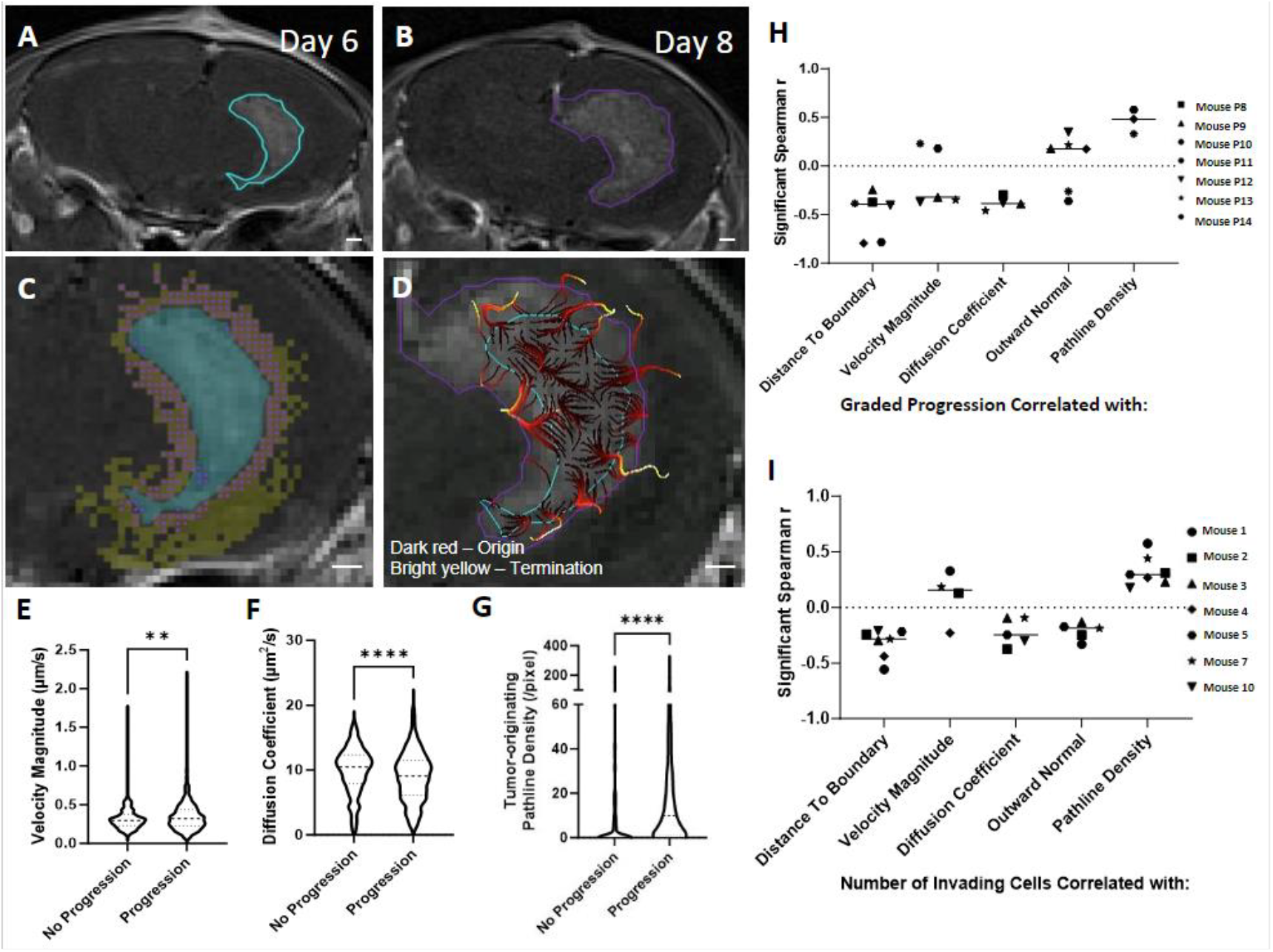
MRI-identified glioma progression correlates with diffusion coefficient, pathlines, and velocity magnitude. A) Day 6 tumor boundary (cyan) as identified on T1 image. B) Day 8 tumor boundary (purple) as identified on a T1 image in the same tumor. C) Pixels containing tumor on Day 8 (purple points) within the contrast-enhancing parenchyma of Day 6 (yellow pixels) are classified as “Progression,” whereas all remaining contrast-enhancing pixels in the parenchyma are classified as “No Progression.” D) Day 8 MRI is overlaid with Day 8 tumor boundary (purple), Day 6 tumor boundary (cyan) and Day 6 tumor-originating pathlines (dark red to yellow transition indicates direction of pathline origination to termination) (all scale bars = 500 μm). E) Local velocity magnitude comparison between no progression and progression regions (Mann-Whitney U test performed, p-value = 0.009; Cohen’s D = 0.15; No Progression (N=435) Progression (N=1524)). F) Local diffusion coefficient comparison between no progression and progression regions (Mann-Whitney U test performed, p-value <0.001; Cohen’s D = 0.26). G) Tumor originating pathline density comparison between no progression and progression regions (Mann-Whitney U test performed, p-value < 0.001; Cohen’s D =0.31). H) Neighborhood multiparametric correlation analysis of MR-signal identified graded progression metric. I) Neighborhood multiparametric correlation analysis of histologically identified invasion. All correlations shown are significant as obtained from a Spearman correlation. Animals not shown for a specific parameter were not significant.

### Both histological invasion and radiographic progression are captured by the same transport patterns

The observed correlations between transport metrics and the spatial radiologic progression mirror the correlations identified in the study where invasion was quantitatively assessed using histological tissue sections (**Figure 6H-I**). Regions of progression and invasion were negatively correlated with distance to boundary, meaning both are occurring close to the tumor boundary, as expected. Regions of progression and invasion were negatively correlated with the diffusion coefficient in four and five of the seven mice, respectively. Tumor-originating pathline density was positively correlated with progression in three of the seven mice and with invasion in all seven mice, indicating regions with increased tumor-originating outward flow can be associated with both progression of disease and invasion. These results indicate that transport metrics can characterize single-cell invasion at a given timepoint while also capturing the broader, cumulative impact of such invasion on disease progression.

## Discussion

In this study, we developed a set of analytical techniques to quantify transport metrics of interstitial fluid flow as determined by non-invasive MR imaging and link those metrics to important physiological outcomes (**Table 1**). We have previously validated the use of DCE-MRI for measuring transport within and around glioma based on underlying physical principles [9]. Our features include distance from delineated tumor boundary, fluid flow velocity, diffusion coefficient, and two novel metrics to quantify fluid moving with respect to the tumor boundary: angle difference of tumor boundary normal vector to fluid flow direction and density of tumor-originating pathlines. We directly linked these transport features to GBM invasion and progression in mice. Specifically, averages of tumor-wide velocity magnitude as well as spatially explicit local analysis of velocity magnitude reveal a positive correlation with invasion. Interestingly, spatial averages of diffusion coefficient exhibit no correlation to invasion, but the diffusion coefficient does have a significant negative correlation to invasion when interrogated at a finer spatial resolution. Notably, tumor-originating pathline density emerges as a novel transport metric associated with invasion. These transport metrics that significantly correlate with histologically identified invading cells are the same metrics that correlate with radiographic progression in our murine xenograft model.

**Table 1:**
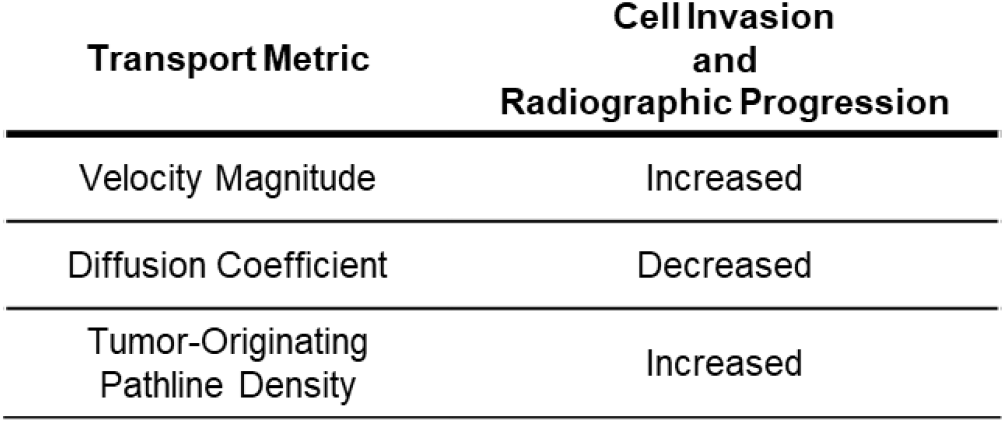
Summary of key dynamics between interstitial fluid flow parameters and glioma cell invasion in murine models.

Elevated velocities have been implicated in a host of cancers, primarily breast and glioma [10], [11], [12], [13]. These elevated velocities have been linked to increased glioblastoma cell invasion in vitro [14] and in vivo [6]. We and others have identified underlying mechanisms that drive flow-induced glioma invasion consisting of both intra- and extracellular drivers [15]. Of interest to the current study, when cells are exposed to subtle interstitial fluid flow, pericellular gradient formations can drive directional migration through autologous chemotaxis [16]. These gradients, as modeled in silico, are sensitive to the features of flow, including magnitudes and directionalities, indicating a potential sensitivity the complex flows that we see through our image analysis [17]. In silico models have explored these complexities that arise due to contributions of the tumor microenvironment, such as heterogeneous vasculature and intratumoral pressure [18], [19]. Seminal work investigating transport in glioma reports intratumoral pressures drive interstitial fluid velocities into the surrounding parenchyma, though the heterogeneity of flow within and around tumors in the brain remained understudied in experimental models [20], [21]. Our localized analysis shows that heterogeneity of fluid flow transport within and around the tumor differentially accounts for spatially varying invasion patterns. Thus, considering local transport trends is important to better understand initial invasion and ultimate progression of the disease.

The diffusion coefficient of the parenchyma has also been associated with invasiveness in cancer. Endometrial, cervical, and breast cancers have a lower diffusion coefficient than comparable healthy tissue [22], [23], [24], [25]. One retrospective clinical study interrogated all voxels as individual data points and, similar to our study, found that regions of progression were most associated with a lower apparent diffusion coefficient [26]. One marked difference between our results and these reports is that we measure the apparent diffusion coefficient of the contrast agent gadolinium through the solution of the mass transport equation. Other studies use the diffusion tensor or apparent diffusion coefficient of water. As the diffusion coefficient is directly related to molecule size, it follows that our calculated rates are slower than these reported rates, due to the difference in size between water and gadolinium. The biological reasons for this relationship are not fully known, as there are seemingly counteracting elements involved. Specifically, diffusion can be limited in denser matrices, with reduced porosity, and thus, cells may have more limited free movement. However, these denser matrices offer binding sites of preferential interactions with receptors on tumor cells that could have an inverse effect on more active motility. Other effects specific to diffusion coefficient relate to the interactions between the substrate measured and the surrounding tissue, such as electrostatic charge, which may also be interacting with cells in interesting ways for further exploration.

As invasion ultimately originates from the tumor bulk and travels into the parenchyma, the ability to examine and quantify downstream implications of tumor-associated drainage is logical but previously unexplored. We developed two novel quantitative markers of the dispersal of fluid outward from the tumor boundary to investigate outward flow. One of these markers, the density of tumor-originating pathlines in the parenchyma, offers knowledge not only about the tumor but also about how the tumor interfaces with the parenchyma and the locations of regions of interest. Locations of elevated pathline accumulations may be local sinks for interstitial fluid flow. Draining vasculature may serve as sinks for gadolinium drainage, specifically funneling into perivascular spaces [27]. These physical factors may be reasons why outward interstitial flow so significantly corresponds to regions of invasion and progression.

The primary aim of the methodology developed here is to apply it to clinical images to gain spatial insight into recurrence and tumor margins in glioma patients. Invasive tumor margins in GBM yield incomplete resection, therapeutic targeting, and, ultimately, recurrence in patients. Thus, the ability to utilize non-invasive standard of care imaging and quantitative metrics like those developed in this study that can aid in the identification of tumor cells at these margins has long been sought [28]. Intraoperative identification of tumor cells has been achieved using imaging agents such as 5-ALA [29] and the field of photodynamic therapies is rapidly improving to incorporate improved imaging agents [30], though these methods are limited to surface identification during surgery, and do not describe further tissue invasion or progression. Methods to leverage MRI combined with AI or machine learning have been probed to define treatment margins or predict progressive disease [31], yet few are grounded in the inherent underlying physical and biological drivers of tumor invasion like fluid transport. Labeling glioma cells with iron-oxide nanoparticles enables tracking of progression along specific structures [32], however cell-labeling methods lack translatability to patients. Clinically relevant approaches like the quantitative analysis developed here must be used to connect patient outcomes with underlying features of the tumor microenvironment.

While we have presented novel quantitative metrics from DCE-MRI, they should not be relied upon in isolation. Our fluid flow metrics can and should be added to the feature space that can be obtained from other MR imaging modalities. Some MRI-calculated features that have been clinically correlated with survival in glioma include: preoperative tumor volume, Karnofsky performance status (KPS), tumor location, edema, extent of resection, contrast enhancement, temporal changes in serum proteomics and pH [33], [34], [35], [36], [37]. Specifically, perfusion imaging has been one of the most clinically used MRI scans in the treatment of glioma providing vessel leakiness from K_Trans_ to identify specific biomarkers indicative of progression and recurrence [38], [39]. These perfusion metrics have also been found to be predictive of tumor grade and treatment response [28]. Combining our novel features with these existing metrics offers a promising approach to improve the accuracy of tumor margin delineation and understanding disease dynamics. For instance, combining these features using more advanced modeling techniques (e.g., linear and non-linear combination classifiers) may yield better performance compared to relying on individual metrics alone.

In conclusion, our study presents a significant stride towards non-invasive, *in vivo* localization of invading tumor cells through the development of quantitative metrics of interstitial fluid flow derived from DCE-MRI. This advancement not only offers the potential to identify additional mechanisms and correlates of invasion via complementary tissue-level analyses such as immunohistochemistry but also opens avenues for exploring extracellular matrix components, metabolites, and cellular reactivities relevant to glioma invasion [40]. Furthermore, a deeper comprehension of transport parameters in and around invasive regions holds promise for the design of novel therapeutic delivery strategies tailored to target these specific locales or capitalize on their unique transport characteristics. Ultimately, this study presents a first step towards the promise of translating interstitial fluid flow analyses into tangible improvements in patient outcomes.

## Materials and Methods

### Cell Culture

GL261-GFP cells were generated as previously described in [12]. Cells were maintained for at least three passages after thawing with Dulbecco’s Modified Eagle Medium (DMEM) + 10% fetal bovine serum (ThermoFisher, Gibco). Cells were resuspended at a concentration of 20,000 cells/uL in serum-free media for tumor implantation as described previously [6]. Glioma stem cells (GSCs) G34 were cultured following the previously described protocol in [41]. GSCs were cultured in neurobasal media (ThermoFisher) with N2 and B27 without vitamin A supplements (ThermoFisher), human recombinant basic fibroblast growth factor (bFGF) and epidermal growth factor (EGF) (50 ng mL−1, ThermoFisher), Glutamax (ThermoFisher), and Penicillin–Streptomycin (ThermoFisher) in low-adhesion tissue culture flasks (Grenier).

### In Vivo Tumor Model

#### Invasion-flow correlation study

For GFP-GL261 injections, all animal procedures were approved by the Institutional Animal Care and Use Committee at Virginia Polytechnic Institute and State University under approved protocol #20-146. Ten, 8–11 month-old transgenic B6.Cg-S1pr3tm1.1Hrose/J mice [42] (JAX stock #028624) were anesthetized with isoflurane and connected to a stereotactic frame. A burr hole was drilled at coordinates −2, +2, −2.2 (AP, ML, DV) from bregma, and 100,000 GFP+ GL261 cells in 5μL serum free DMEM were injected via a Hamilton syringe and syringe pump (World Precision Instruments) at 1μL/min for 5 min. The syringe was inserted to a depth of 3 mm then retracted 0.5 mm before injection. The Hamilton syringe was removed three minutes after the completion of injection to prevent reflux, and the burr hole was sealed with bone wax.

#### Progression study

The progression study was conducted at the University of Virginia under approved protocol 4021. Seven 8–10-week-old SCID mice were injected using the same protocol as previously published in [9]. Mice were injected with G34 GSCs, and the tumors were allowed to grow for 10 days. T1, T2, and DCE-MRI imaging were performed on days 6, 8, and 10. Following DCE-MRI, mice were euthanized via cardiac perfusion, and brains were harvested.

#### Magnetic Resonance Imaging

#### Invasion-flow correlation study

DCE-MRI was performed on all mice with detectable tumors on Days 15 and 24 to allow for tumor growth. Mice were imaged with a 9.4 T small animal MRI (Bruker BioSpec AVANCE NEO 94/20 USR, Ettlingen, Germany) equipped with 660 mT/m high power gradient. An active detunable 86 mm volume coil was used as the transmit coil and a planar 20 mm receive-only surface coil was used as the receive coil. A T2-weighted image was taken using rapid acquisition with relaxation enhancement (RARE) sequence to confirm tumor growth with the following parameters: repetition time (TR) = 1800 ms, echo time (TE) = 40ms, field of view (FOV) = 19.2 × 19.2 mm with a 192 × 192 matrix, slice thickness = 0.5 mm, number of slices = 16, with 9 averages requiring a total acquisition time of about 6.5 min per mouse.

To measure the interstitial fluid transport metrics, a pre-contrast T1-weighted image was collected followed by tail vein injection of Gadolinium (Magnevist, Bayer HealthCare Pharmaceuticals) at a concentration of 0.1mmol/kg in sterile, heparinized saline. Four post-contrast T1-weighted images were acquired for approximately 12 min post-injection, as previously published [4]. T1-weighted images were acquired using a fast low angle shot (FLASH) sequence using the following parameters: repetition time (TR) = 180 ms, echo time (TE) = 11 ms, field of view (FOV) = 19.2×19.2 mm with a 192 × 192 matrix, slice thickness = 0.5 mm, number of slices = 16, and 7 averages requiring a total acquisition time of about 3 min per sequence (**Supplemental Figure 1**).

#### Progression Study

For GSC-injected mice, DCE-MRI was performed on days 6, 8, and 10 according to the protocol established in prior publications [4], [9].

### Tissue Harvest, Cryosectioning, Immunohistochemistry (IHC), and Microscopy

Mice were euthanized and transcardially perfused with ice cold 1 x phosphate buffered saline (PBS) followed by 4% paraformaldehyde (PFA). Brains were post-fixed in PFA for 18 hours and placed in 30% sucrose until complete submersion was achieved. Afterwards, brains were placed in molds of Optical Cutting Temperature (OCT) compound at -80°C and sectioned at 12 um thickness on a cryostat. Specific MRI slices of interest were identified on T2-weighted MRI based on tumor size and minimal needle-track damage. One to two distinct MRI tumor locations were sectioned per mouse. Structural features (i.e., ventricles, white matter, tumor shape) in the T2-weighted MRI images were used as a guide during cryosectioning to collect cryosectioned slices corresponding to the identified MRI slice of interest. At each location, two to three histological sections were used for the analysis based on tissue quality (**Supplemental Figure 2)**. Slides were stained with DAPI (Thermofisher) for 10 minutes and imaged at 20X on a VS200 Olympus Slide Scanner with DAPI in the blue channel and GFP+ GL261 cells in the green channel.

### Registration of MRI Slices to Histological Tissue Sections

Control point image registration was performed in MATLAB. A local weighted mean geometric transformation was used to account for orientation and shape differences between the MRI slices and histological tissue section. At least 12 distinct, reliable anatomical landmarks in both the MRI and histological tissue sections were chosen for each registration. Registration control points were selected on gross anatomy such as the surrounding skull and ventricles. IHC-identified spatial coordinates of the invading GL261 cells and vertices of the tumor boundaries were registered to the MRI using the resulting transformation matrix. A composite tumor boundary was created for each MRI image that preserves only areas identified as tumor bulk in all histological technical replicates (**Supplemental Figure 3**).

### Identification of Tumor Boundary and Invading Tumor Cells

The DAPI and GFP channels were split and exported individually using ImageJ. The green channel of the GFP+ GL261 cells were thresholded based on intensity to remove background fluorescence. Images were colocalized with DAPI staining to include only the GFP signal associated with a nucleus (DAPI). ImageJ watershed segmentation was used to separate touching or overlapping nuclei. The resulting identified objects represented individual GL261 cells.

In this study, invading cells were defined as those no longer adjoined to the contiguous tumor bulk. Two investigators (to improve the rigor and confidence of invading cell selection) worked together to categorize each cell in all histological tissue sections as bulk or invading. To establish a tumor boundary, the (x,y) coordinates of the GFP+ cells of the main tumor bulk were provided as input to the MATLAB *boundary* function. This function created a bounding area that enveloped these points and returned a set of (x,y) vertices of the tumor boundary (**Supplemental Figure 4**).

### Transport Metric Analysis

From the DCE-MRI sequence, transport metrics were obtained using “Lymph4D,” a previously published and openly available tool [8]. Briefly, parameter optimization is performed in a pixel-wise manner for the two-dimensional (2D) diffusion-advection equation to identify the combination of the velocity magnitude and direction of advection and diffusion coefficient that best describes the change in gadolinium over time of that pixel acquired by the DCE-MRI.

### Defining the Contrast-Enhancing Parenchymal Region

To ensure that transport metrics were obtained in pixels that experienced influx of gadolinium, we developed a strategy to define these regions. To determine whether a pixel experienced gadolinium signal or general MRI noise, we used an analysis of the standard deviation of the time-signal intensity curve (TSIC) of each pixel. The standard deviation of the TSIC for each pixel is calculated using the following formula:

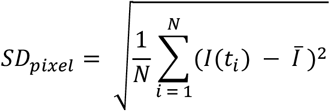

where N is the number of time points in the DCE-MRI, I(t_i_) is the intensity of the pixel at time t_i_, and Īis the mean intensity of the TSIC of that pixel over all time points. A region growing algorithm is then used to semi-automatically identify pixels with standard deviation (SD) of TSIC relatively similar to the tumor bulk. First, a seed pixel was manually selected within the tumor bulk. A region was then iteratively expanded from this pixel by comparing the SD of the neighboring pixels to the mean of the region’s SD. The neighboring pixel with the smallest difference was allocated to the region. This iterative process was continued until the SD difference between the region’s mean and the neighboring pixels becomes larger than a threshold. This threshold was maximized for each DCE-MRI individually to include the maximum number of pixels in the region without negating the iterative process and including every pixel in the image in the region. The result was a semi-automatically identified region of contrast-enhanced parenchyma where the transport metrics calculated using Lymph4D were considered valid and not noise (**Supplemental Figure 4**).

### Definition of Outward Flow Metrics

*Flow normal to boundary* is defined as the difference between the IFF direction vector at each pixel and normal vector at the nearest point on the tumor boundary. For a given pixel, the closest point to the tumor boundary was identified by finding the boundary vertex with the minimum Euclidean distance to the center of the pixel. To calculate the normal vector at this boundary vertex, two lines extending out from this boundary vertex were identified using the two adjacent boundary vertices. The normal vectors of these lines are the perpendicular lines that point away from the tumor bulk. The average direction of these two normal vectors were weighted by the distance away from the boundary vertex of interest to give a more accurate estimate of the normal vector at the boundary vertex of interest. The flow difference to outward normal metric was calculated by taking the angle difference between the IFF direction vector and this boundary normal vector. This metric has a value between [0, 180], where zero indicates an IFF flow pointing directly away from the tumor boundary and 180 indicates an IFF flow direction pointing directly toward the tumor boundary.

*Tumor-originating pathlines* were obtained by tracking the trajectories of virtual particles with origination points within the tumor boundary. A pathline is obtained by seeding a particle at a location given by 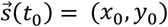 within the MRI image. The virtual particle seeded at this location will have a velocity given by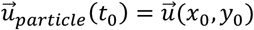where 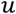is defined by the velocity of the flow field provided by DCE-MRI analysis. The particle trajectory 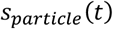 is calculated using ODE45 in MATLAB. This method places a particle at the center of each pixel contained within the previously calculated tumor boundary and calculates the trajectory until termination, meaning it reaches a velocity equal to zero. For a given pixel, the number of tumor-originating pathlines that traverse the pixel is summed and reported as the “tumor-originating pathline density.”

### Tumor- and Contrast-Enhanced Parenchymal-Wide Analysis

Each transport metric was averaged for the overall tumor area and for the contrast-enhanced parenchymal region. *Statistical analysis:* Unpaired Mann-Whitney U tests and Spearman r correlations were used to compare averaged tumor and contrast-enhanced parenchymal transport metrics.

### Local Metric Analysis

For each pixel within the tumor boundary and the identified contrast-enhanced parenchyma, a feature vector was constructed including the following metrics: distance to boundary, flow magnitude, diffusion coefficient, flow direction angle difference to boundary normal, tumor-originating pathline density, and number of invading cells. The distance to boundary metric is calculated as the Euclidean distance between the focal center of the pixel and the nearest point on the tumor boundary. The flow magnitude, diffusion coefficient, and flow direction angle difference to outward normal are calculated by averaging this value of each metric in the 3×3 pixel neighborhood surrounding the pixel. For the tumor-originating pathline density and the number of invading cells, the values of these metrics in all nine individual pixels in the 3×3 pixel neighborhood are summed (**Supplemental Dataset 1**). *Statistical analysis:* Unpaired Mann-Whitney t-tests were used to compare transport metrics in pixels with and without invasion. Pearson r correlations were used for multiparametric correlation analysis.

### Progression Analysis on Tumor Growth Differences

For each mouse, a T1-weighted MRI slice of interest was manually identified that included the largest tumor area observed six days post tumor injection. IFF transport metrics were calculated, and contrast-enhanced parenchyma was identified using the DCE-MRI of this slice of interest. Manual slice matching of T2-weighted images to the DCE-MRI modality was required for tumor segmentation. Manual slice matching was performed by concurrently comparing the lesion and surrounding anatomy between the DCE-MRI and T2-weighted images from both six- and eight-days post injection. Anatomically matched T2-weighted images from six- and eight-days post injection were registered to the T1-weighted slice of interest using control point registration with a nonreflective similarity transformation in MATLAB. On these registered T2-weighted images, the tumor was manually segmented, and the regions segmented as tumor eight days post injection that were not segmented as tumor six days post injection were identified as tumor progression. Any pixel on the T1-weighted slice of interest, including any part of this identified progression region, was classified as a progression pixel. “Graded progression is defined as the number of pixels in the 3×3 pixel neighborhood classified as progression, with possible values from [0,9]. *Statistical analysis:* Unpaired Mann-Whitney t-tests were used to compare transport metrics in pixels with and without progression. Pearson r correlations were used for multiparametric correlation analysis.

## Acknowledgements

The authors would like to thank Maosen Wang, Jack Roy, and Jeremy Gatesman for assistance with MRI, Samantha Haynes, Ehaab Saleem, Micaiah Lee, and Skylar Davis, for assistance with MRI image analysis and tissue sectioning. This work was funded by NCI R37CA222563 to JMM, the RedGates Foundation to JMM (Virginia Tech), American Cancer Society Institutional Research Grant to JMM (UVa), and NINDS R01NS115971 to RCR and JMM, NSF GRFP to KMK and CMC.

## Supplemental Data and Figures

**Supplemental Figure 1:** DCE-MRI sequence of contrast agent movement.

**Supplemental Figure 2:** Sampling and matching of histological slides and MRI.

**Supplemental Figure 3:** MRI and IHC section matching and control point registration.

**Supplemental Figure 4**: Invading cell identification and post registration data.

**Supplemental Figure 5:** GL261 tumor velocity and diffusion variability.

**Supplemental Dataset 1 – Image analysis on implanted GL261 tumors** (A) Tumor and contrast-enhanced parenchyma converted to masked regions overlaid a T1-weighted MRI. (B) Tumor and contrast-enhanced parenchyma converted to masked regions overlaid a T1-weighted MRI with invading tumor cells. (C) Heat map of velocity magnitude on the whole brain with tumor and contrast-enhanced parenchyma outlined. (D) Heat map of diffusion coefficient with the tumor and contrast-enhanced parenchyma outlined. (E) Tumor-originating pathlines overlaid on T1-weighted contrast-enhanced MRI with tumor and contrast-enhanced parenchyma. (F) Tumor-originating pathline density for each pixel with tumor and contrast-enhanced parenchyma. Dark blue and dark red represent a low and high tumor-originating pathline density, respectively.

**Supplemental Dataset 2 – Image analysis on implanted G34 tumors with progression** (A) Day 6 tumor boundary as identified on T1-weighted contrast enhanced MR image. (B) Day 8 tumor boundary as identified on a T1-contrast-enhanced MR image in the same tumor. (C) Pixels displaying progression within the contrast-enhancing parenchyma identified by Day 6 DCE-MRI. Pixels containing tumor on Day 8 within the contrast-enhancing parenchyma of Day 6 are classified as “Progression,” whereas all remaining contrast-enhancing pixels in the parenchyma are classified as “No Progression.” (D) Day 8 MRI is overlaid with Day 8 tumor boundary, Day 6 tumor boundary and Day 6 tumor-originating pathlines.

