## Supplemental Tables and Figures for "Interstitial fluid transport dynamics predict glioblastoma invasion and progression"

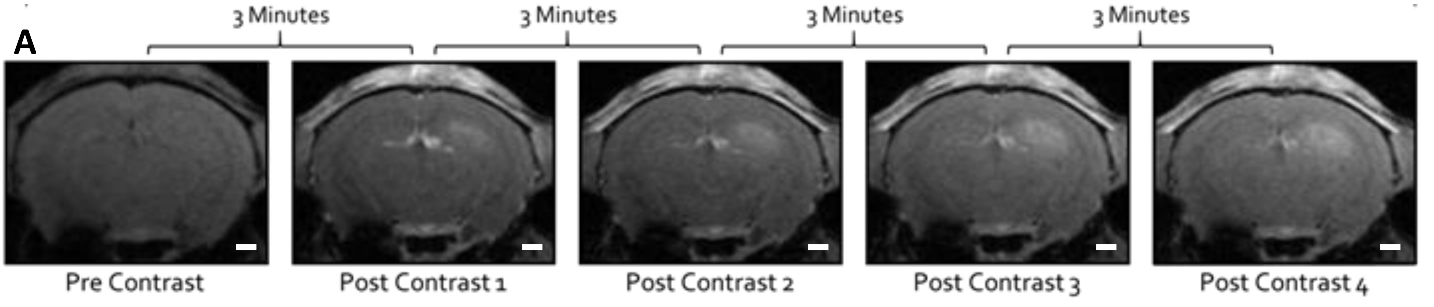


**Supplemental Figure 1: DCE-MRI sequence of contrast agent movement.** A pre-contrast image is acquired followed by four post-contrast images for a total acquisition time of ~12mins (Scale bar = 1mm).

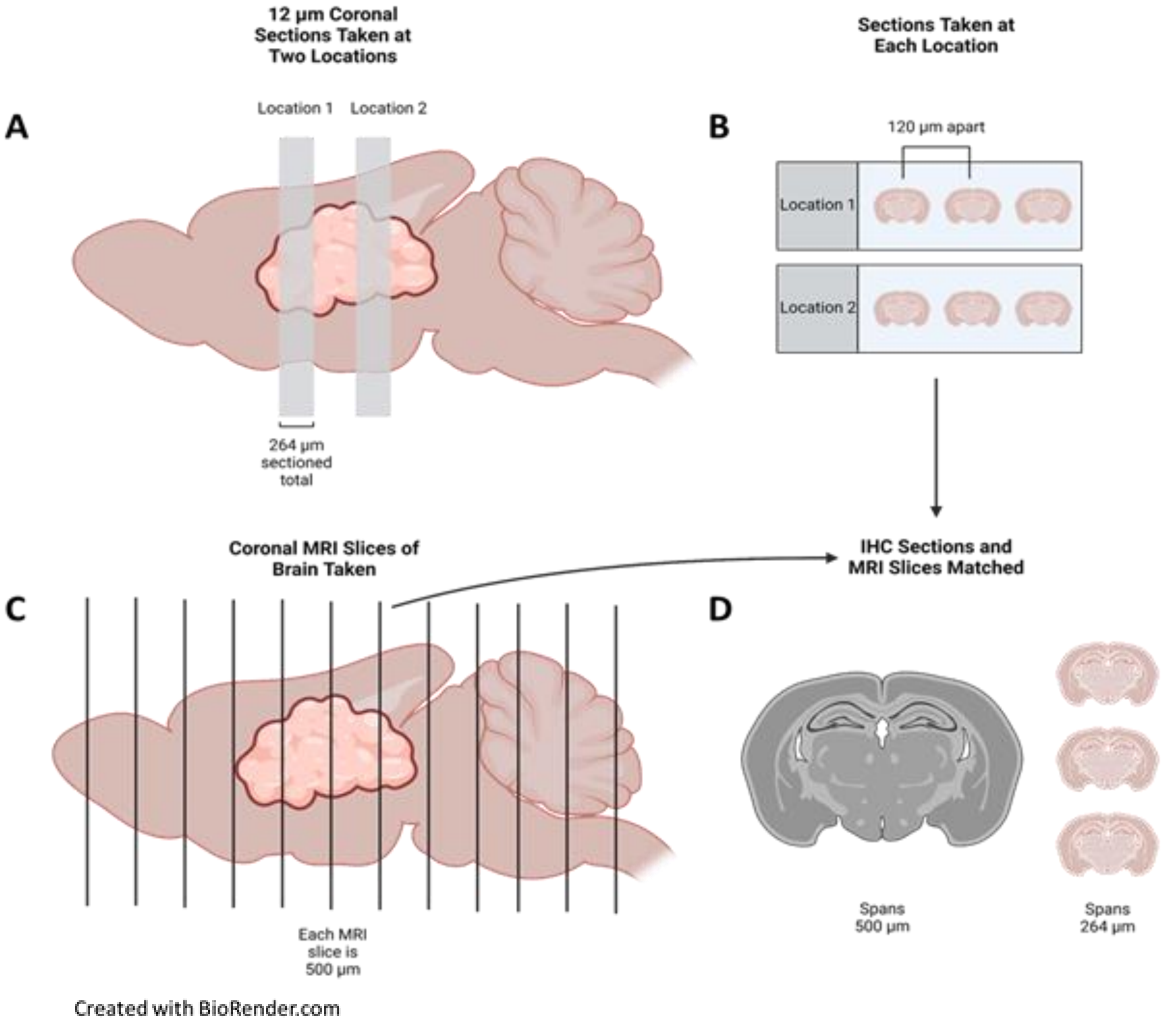


**Supplemental Figure 2: Sampling and matching of histological slides and MRI.** (A) Two distinct locations of the ex vivo mouse brain and tumor are sectioned in 12µm sections for a total of 264µ m at each location. (B) Slides are created with 2-3 brain sections per slide that are 120 µm apart. (C) MRI is performed on the whole mouse brain with each section being 500 µm thick. (D) The tissue sections are matched to the corresponding MRI slice so that the 2-3 sections fall within the 500µm MRI slice.

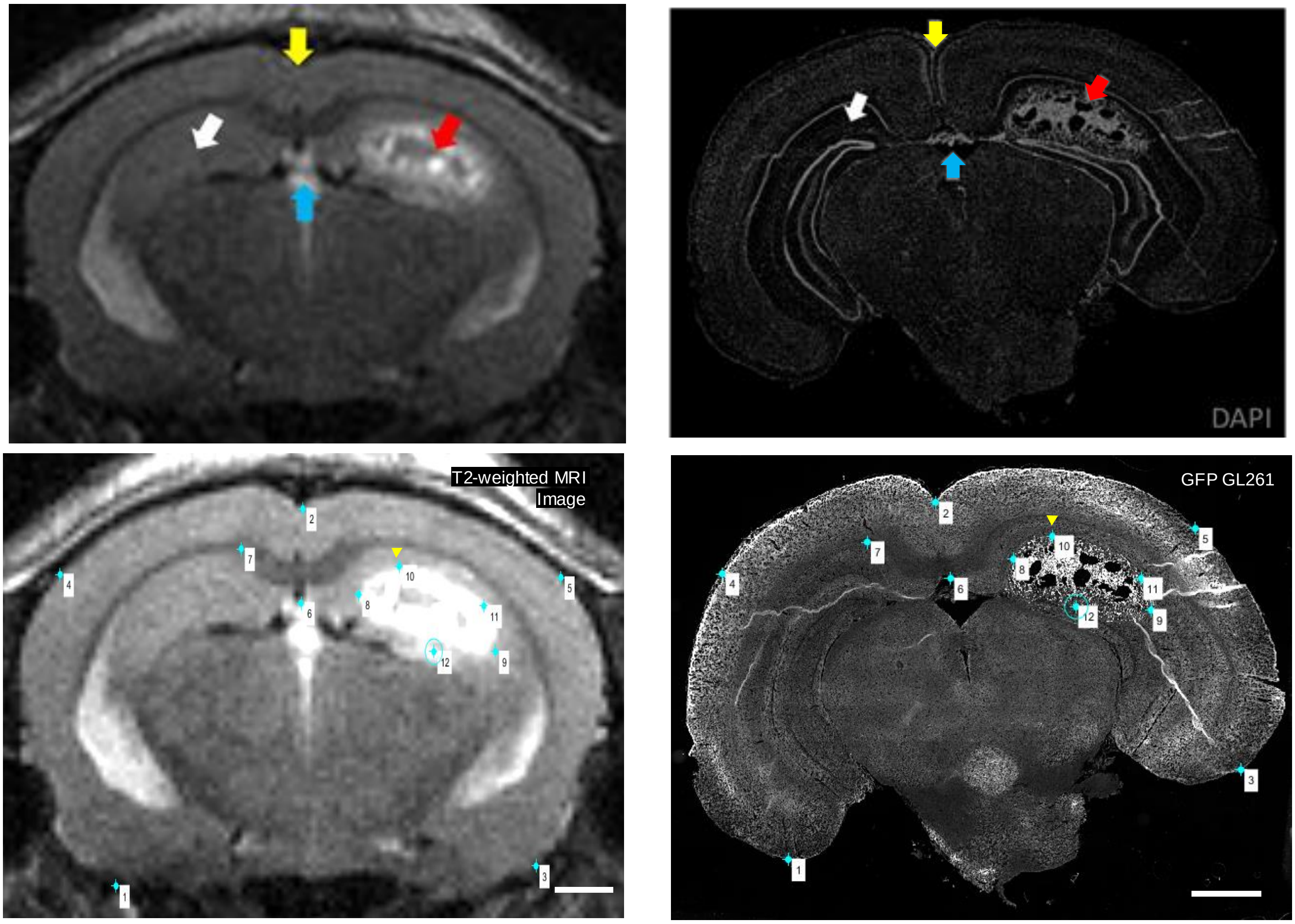


**Supplemental Figure 3: MRI and IHC section matching and control point registration.** MRI slice showing enhanced tumor region (red arrow), midline (yellow arrow), hippocampal region (white arrow), and third ventricle (blue arrow) alongside IHC section stained with DAPI showing tumor border (red arrow), midline (yellow arrow), hippocampal region (white arrow) and third ventricle (blue arrow). 12 control points are chosen on the T2-weighted MRI and GFP GL261 IHC slice for control point registration.

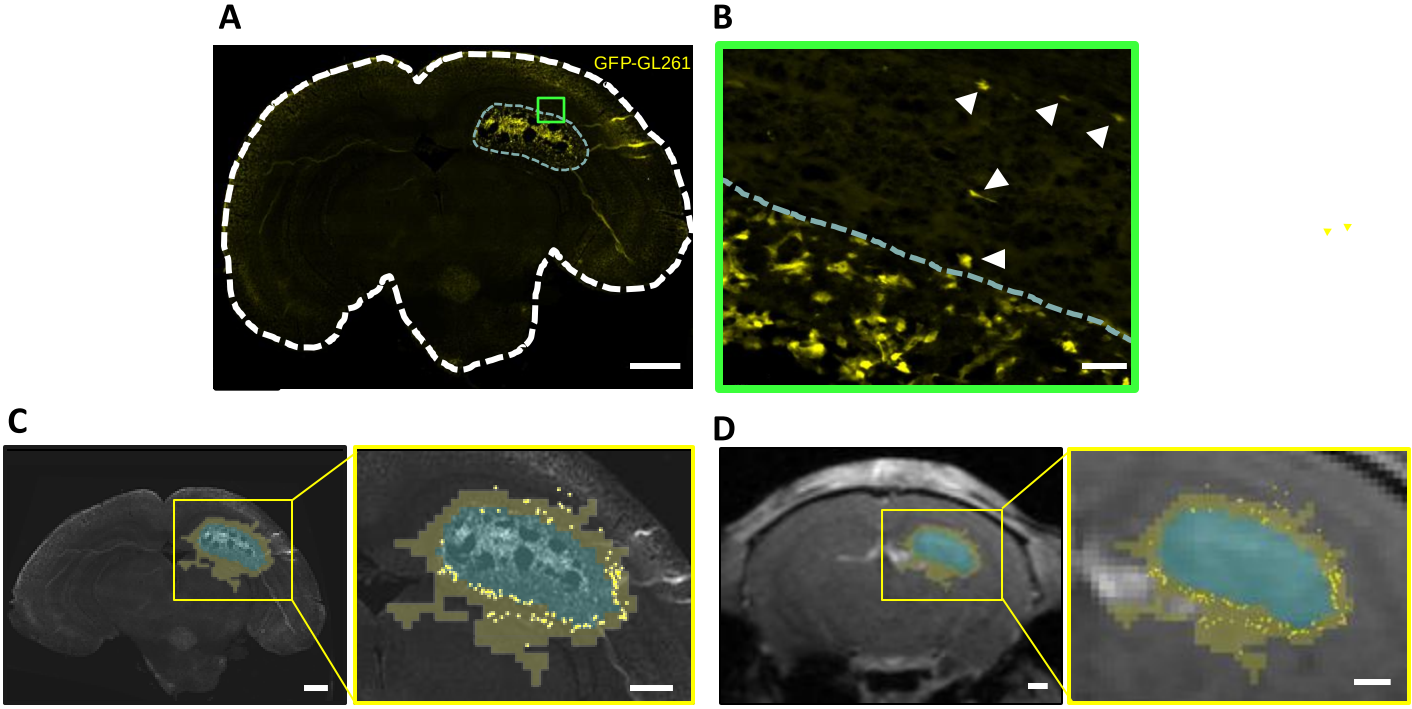


**Supplemental Figure 4: Invading cell identification and post registration data.** A) GFP-GL261 tumor cells (green) and the tumor border (dotted white line) were identified and delineated on histology. B) Arrowheads represent invading cells (scale bar large image = 800µm; scale bar inset = 50µm). C) Tumor (light blue highlight) and contrast-enhanced parenchyma (light yellow highlight) converted to masked regions overlaid with invading tumor cells (dark yellow points) overlaid on fluorescently labeled GFP-GL261 histology and a T2-weighted MRI (D) (Scale bar =1mm, scale bar inset = 500µm) .

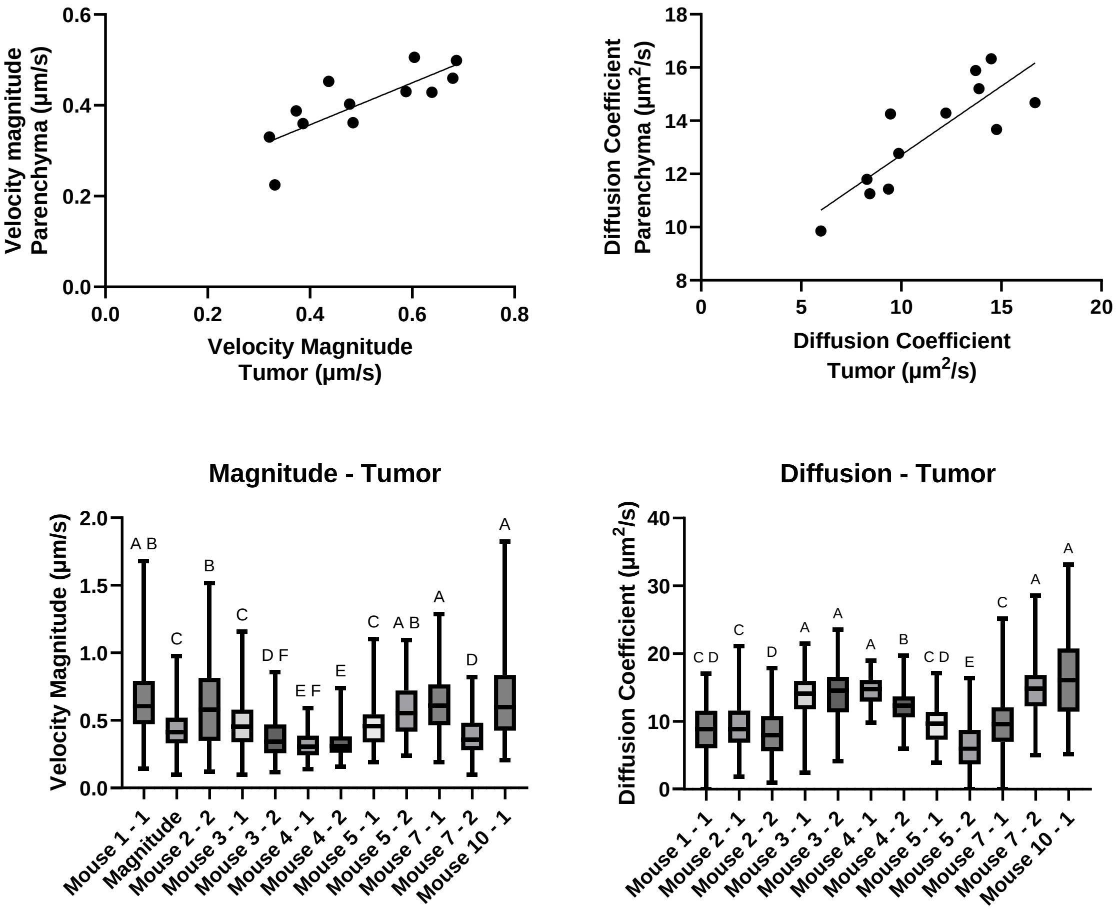


**Supplemental Figure 5: GL261 tumor velocity and diffusion variability.** The average velocity magnitudes within the tumor vs the surrounding parenchyma are positively correlated, meaning that mice with relatively faster velocities in the tumor also have faster velocities in the parenchyma. Diffusion coefficients in the tumor and the contrast-enhanced parenchyma were significantly positively correlated (Spearman r = 0.7902, p-value = 0.0033). The velocity magnitude and diffusion coefficient within the tumor for all individual mouse locations. Compact letter display shows columns are statistically indistinguishable if and only if they share at least one letter.
