## Supplemental Dataset 1 for "Interstitial fluid transport dynamics predict glioblastoma invasion and progression"

### Supplemental Dataset 1: Image analysis on implanted GL261 tumors

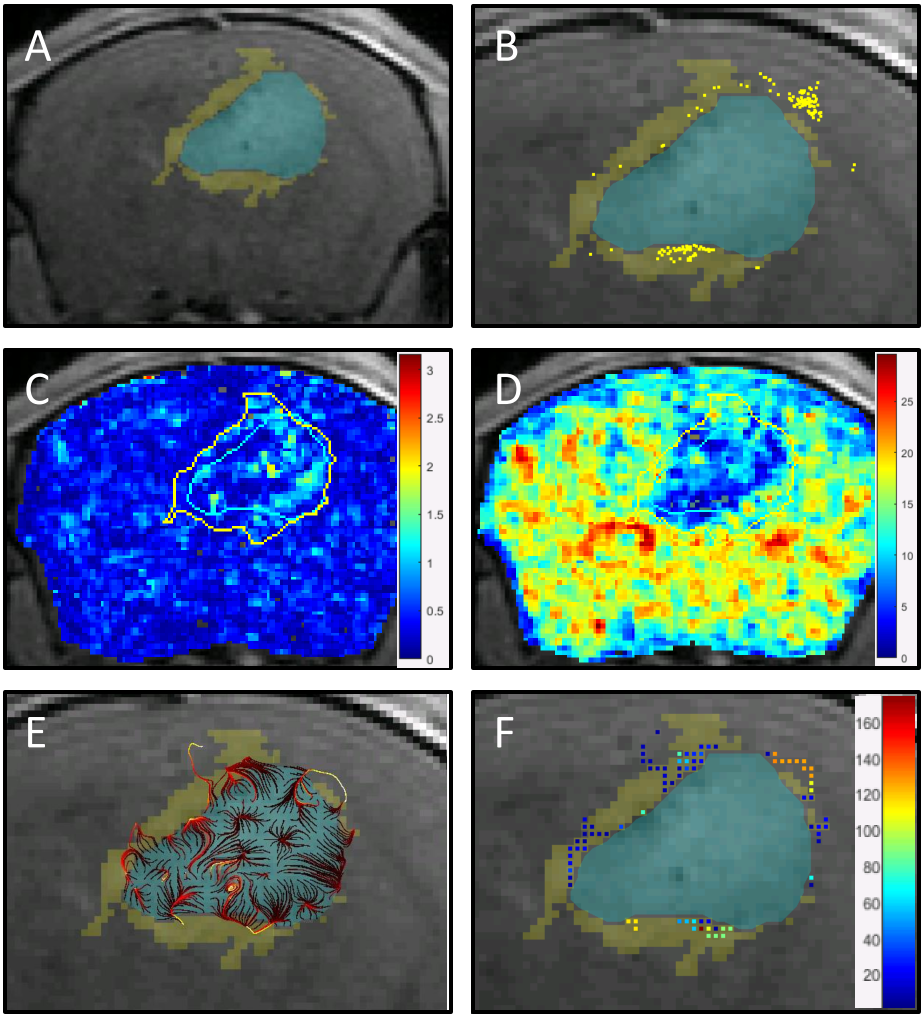


**Supplemental Dataset 1 – GL261 tumors - Mouse 2 – Location 1.** (A) Tumor (light blue highlight) and contrast-enhanced parenchyma (light yellow highlight) converted to masked regions overlaid a T1-weighted MRI (Scale bar =1mm) (B) Tumor (light blue highlight) and contrast-enhanced parenchyma (light yellow highlight) converted to masked regions overlaid a T1-weighted MRI with invading tumor cells (dark yellow points) (Scale bar = 500µm). C) Heat map of velocity magnitude on the whole brain with tumor (light blue) and contrast-enhanced parenchyma(yellow) outlined (Scale bar =1mm). D) Heat map of diffusion coefficient with the tumor (light blue) and contrast-enhanced parenchyma (yellow) outlined (Scale bar =1mm). E) Tumor-originating pathlines (dark red is initial location moving to ending location as bright yellow) overlaid on T1-weighted contrast-enhanced MRI with tumor (light blue highlight) and contrast-enhanced parenchyma (light yellow highlight) (Scale bar =500µm). F) Tumor-originating pathline density for each pixel with tumor (light blue highlight) and contrast-enhanced parenchyma (light yellow highlight). Dark blue and dark red represent a low and high tumor-originating pathline density, respectively (Scale bar =500µm).

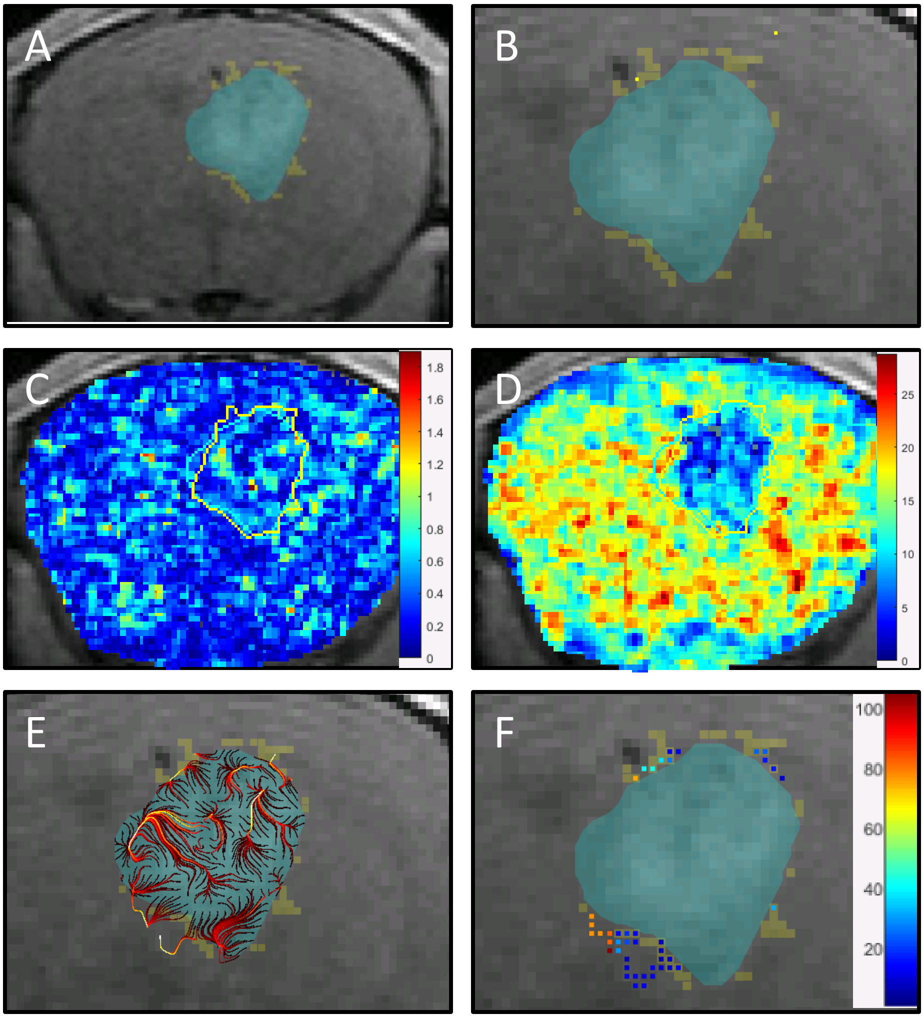


**Supplemental Dataset 1 – GL261 tumors - Mouse 2 – Location 2.** (A) Tumor (light blue highlight) and contrast-enhanced parenchyma (light yellow highlight) converted to masked regions overlaid a T1-weighted MRI (Scale bar =1mm) (B) Tumor (light blue highlight) and contrast-enhanced parenchyma (light yellow highlight) converted to masked regions overlaid a T1-weighted MRI with invading tumor cells (dark yellow points) (Scale bar = 500µm). C) Heat map of velocity magnitude on the whole brain with tumor (light blue) and contrast-enhanced parenchyma(yellow) outlined (Scale bar =1mm). D) Heat map of diffusion coefficient with the tumor (light blue) and contrast-enhanced parenchyma (yellow) outlined (Scale bar =1mm). E) Tumor-originating pathlines (dark red is initial location moving to ending location as bright yellow) overlaid on T1-weighted contrast-enhanced MRI with tumor (light blue highlight) and contrast-enhanced parenchyma (light yellow highlight) (Scale bar =500µm). F) Tumor-originating pathline density for each pixel with tumor (light blue highlight) and contrast-enhanced parenchyma (light yellow highlight). Dark blue and dark red represent a low and high tumor-originating pathline density, respectively (Scale bar =500µm).

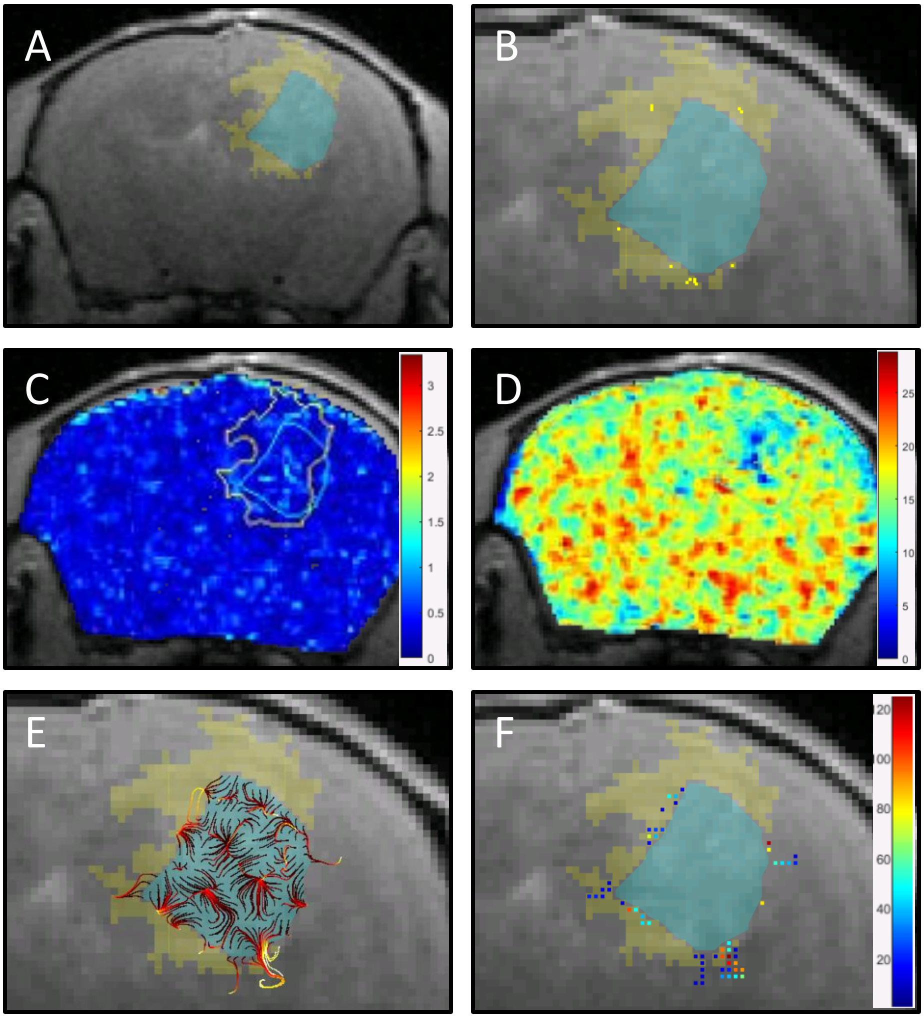


**Supplemental Dataset 1 – GL261 tumors - Mouse 3 – Location 1.** (A) Tumor (light blue highlight) and contrast-enhanced parenchyma (light yellow highlight) converted to masked regions overlaid a T1-weighted MRI (Scale bar =1mm) (B) Tumor (light blue highlight) and contrast-enhanced parenchyma (light yellow highlight) converted to masked regions overlaid a T1-weighted MRI with invading tumor cells (dark yellow points) (Scale bar = 500μm). C) Heat map of velocity magnitude on the whole brain with tumor (light blue) and contrast-enhanced parenchyma(yellow) outlined (Scale bar =1mm). D) Heat map of diffusion coefficient with the tumor (light blue) and contrast-enhanced parenchyma (yellow) outlined (Scale bar =1mm). E) Tumor-originating pathlines (dark red is initial location moving to ending location as bright yellow) overlaid on T1-weighted contrast-enhanced MRI with tumor (light blue highlight) and contrast-enhanced parenchyma (light yellow highlight) (Scale bar =500μm). F) Tumor-originating pathline density for each pixel with tumor (light blue highlight) and contrast-enhanced parenchyma (light yellow highlight). Dark blue and dark red represent a low and high tumor-originating pathline density, respectively (Scale bar =500μm).

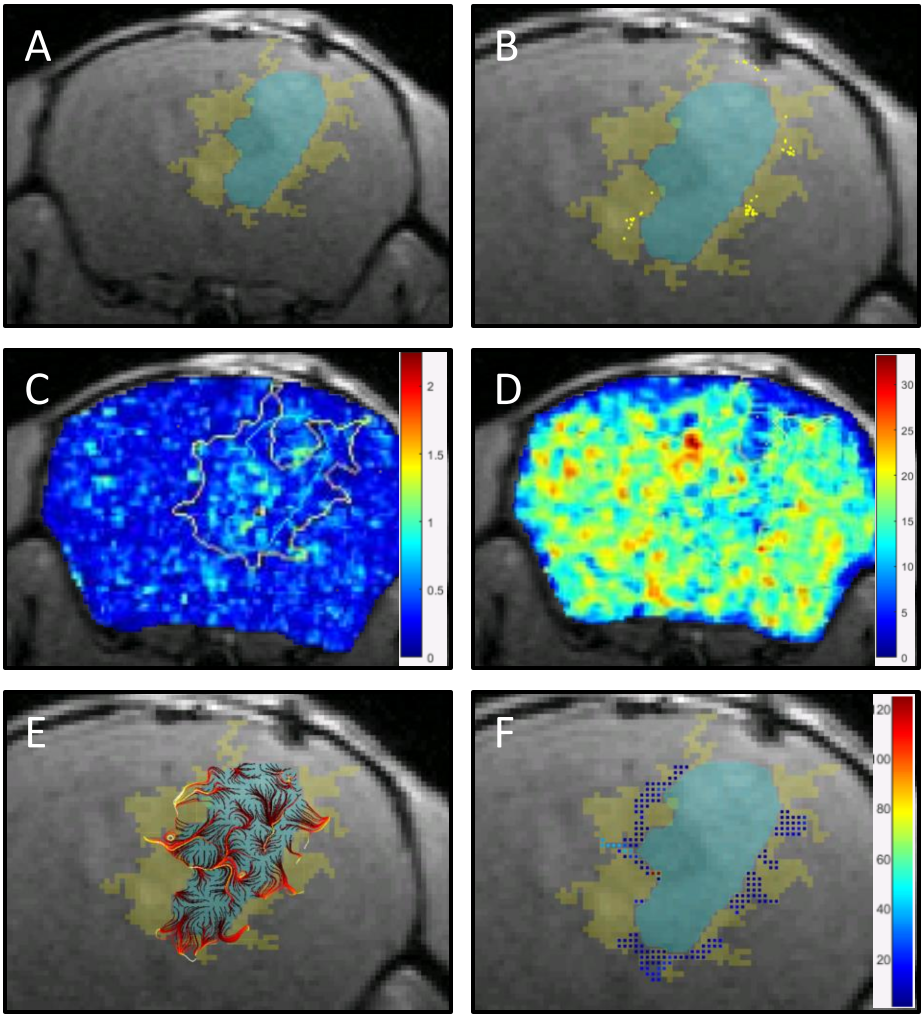


**Supplemental Dataset 1 – GL261 tumors - Mouse 3 – Location 2.** (A) Tumor (light blue highlight) and contrast-enhanced parenchyma (light yellow highlight) converted to masked regions overlaid a T1-weighted MRI (Scale bar =1mm) (B) Tumor (light blue highlight) and contrast-enhanced parenchyma (light yellow highlight) converted to masked regions overlaid a T1-weighted MRI with invading tumor cells (dark yellow points) (Scale bar = 500µm). C) Heat map of velocity magnitude on the whole brain with tumor (light blue) and contrast-enhanced parenchyma(yellow) outlined (Scale bar =1mm). D) Heat map of diffusion coefficient with the tumor (light blue) and contrast-enhanced parenchyma (yellow) outlined (Scale bar =1mm). E) Tumor-originating pathlines (dark red is initial location moving to ending location as bright yellow) overlaid on T1-weighted contrast-enhanced MRI with tumor (light blue highlight) and contrast-enhanced parenchyma (light yellow highlight) (Scale bar =500µm). F) Tumor-originating pathline density for each pixel with tumor (light blue highlight) and contrast-enhanced parenchyma (light yellow highlight). Dark blue and dark red represent a low and high tumor-originating pathline density, respectively (Scale bar =500µm).

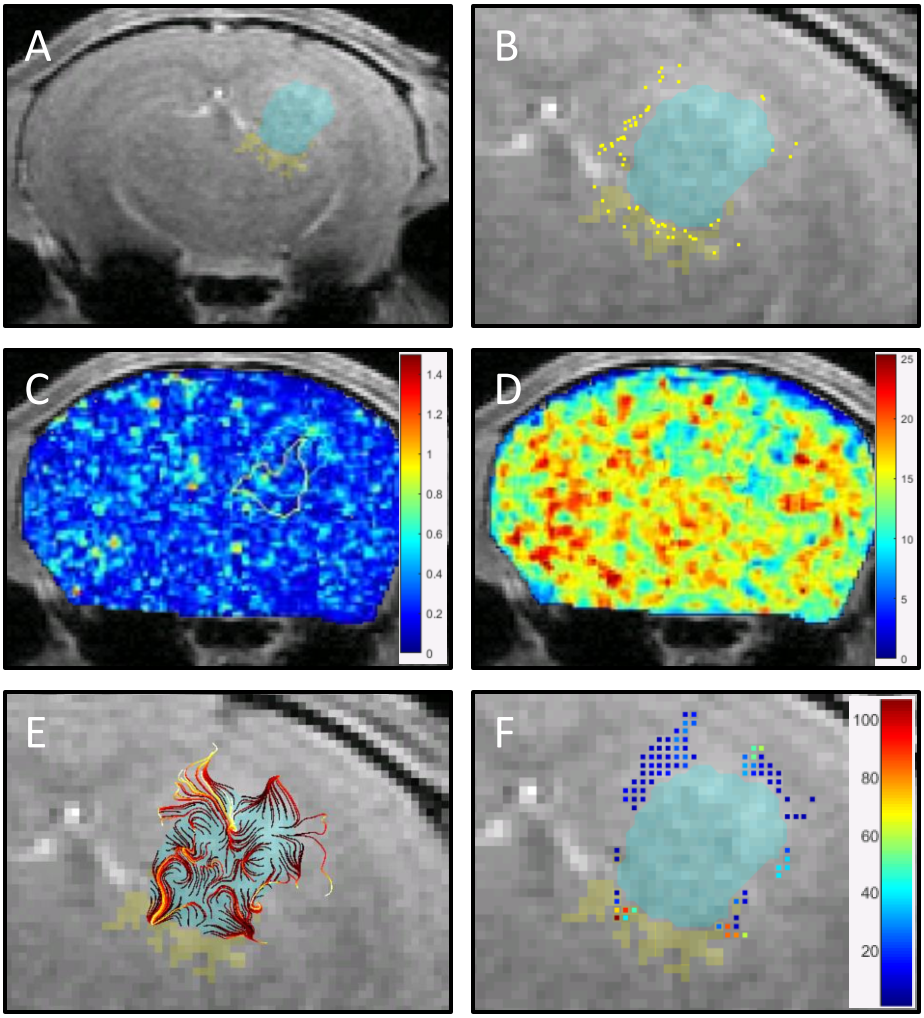


**Supplemental Dataset 1 – GL261 tumors - Mouse 4 – Location 1.** (A) Tumor (light blue highlight) and contrast-enhanced parenchyma (light yellow highlight) converted to masked regions overlaid a T1-weighted MRI (Scale bar =1mm) (B) Tumor (light blue highlight) and contrast-enhanced parenchyma (light yellow highlight) converted to masked regions overlaid a T1-weighted MRI with invading tumor cells (dark yellow points) (Scale bar = 500µm). C) Heat map of velocity magnitude on the whole brain with tumor (light blue) and contrast-enhanced parenchyma(yellow) outlined (Scale bar =1mm). D) Heat map of diffusion coefficient with the tumor (light blue) and contrast-enhanced parenchyma (yellow) outlined (Scale bar =1mm). E) Tumor-originating pathlines (dark red is initial location moving to ending location as bright yellow) overlaid on T1-weighted contrast-enhanced MRI with tumor (light blue highlight) and contrast-enhanced parenchyma (light yellow highlight) (Scale bar =500µm). F) Tumor-originating pathline density for each pixel with tumor (light blue highlight) and contrast-enhanced parenchyma (light yellow highlight). Dark blue and dark red represent a low and high tumor-originating pathline density, respectively (Scale bar =500µm).

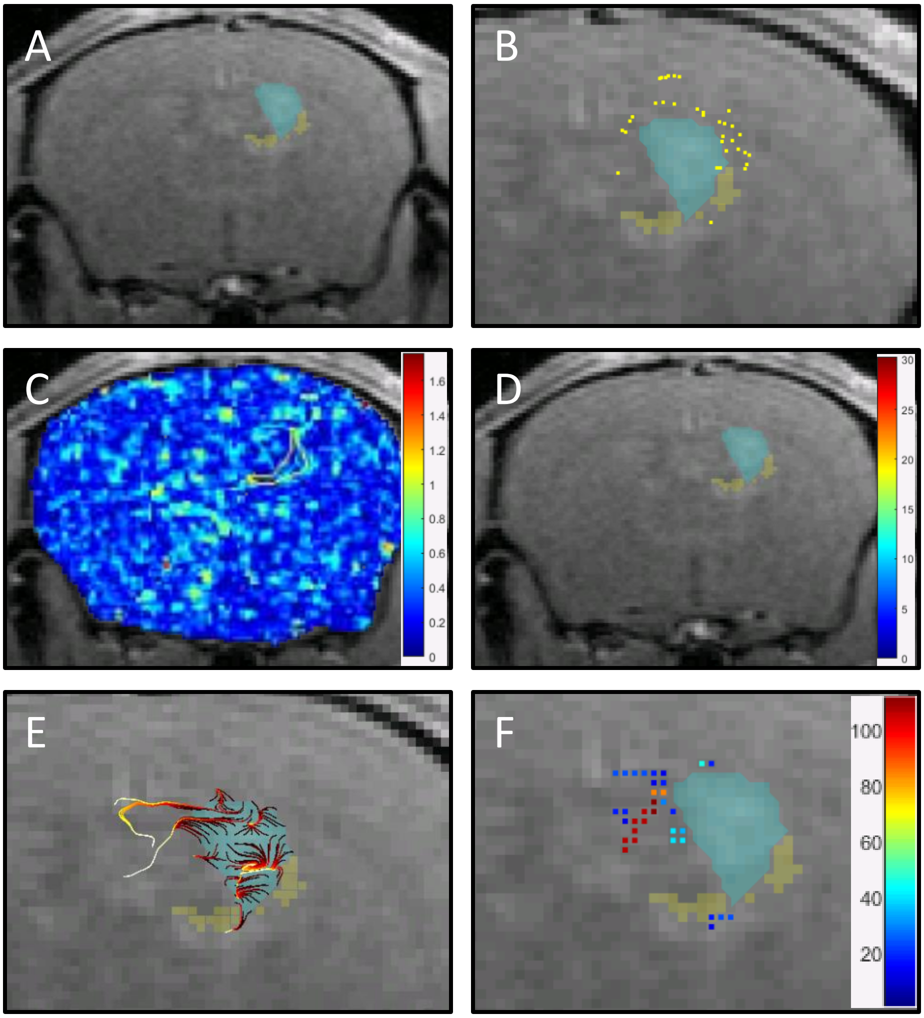


**Supplemental Dataset 1 – GL261 tumors - Mouse 4 – Location 2.** (A) Tumor (light blue highlight) and contrast-enhanced parenchyma (light yellow highlight) converted to masked regions overlaid a T1-weighted MRI (Scale bar =1mm) (B) Tumor (light blue highlight) and contrast-enhanced parenchyma (light yellow highlight) converted to masked regions overlaid a T1-weighted MRI with invading tumor cells (dark yellow points) (Scale bar = 500µm). C) Heat map of velocity magnitude on the whole brain with tumor (light blue) and contrast-enhanced parenchyma(yellow) outlined (Scale bar =1mm). D) Heat map of diffusion coefficient with the tumor (light blue) and contrast-enhanced parenchyma (yellow) outlined (Scale bar =1mm). E) Tumor-originating pathlines (dark red is initial location moving to ending location as bright yellow) overlaid on T1-weighted contrast-enhanced MRI with tumor (light blue highlight) and contrast-enhanced parenchyma (light yellow highlight) (Scale bar =500µm). F) Tumor-originating pathline density for each pixel with tumor (light blue highlight) and contrast-enhanced parenchyma (light yellow highlight). Dark blue and dark red represent a low and high tumor-originating pathline density, respectively (Scale bar =500µm).

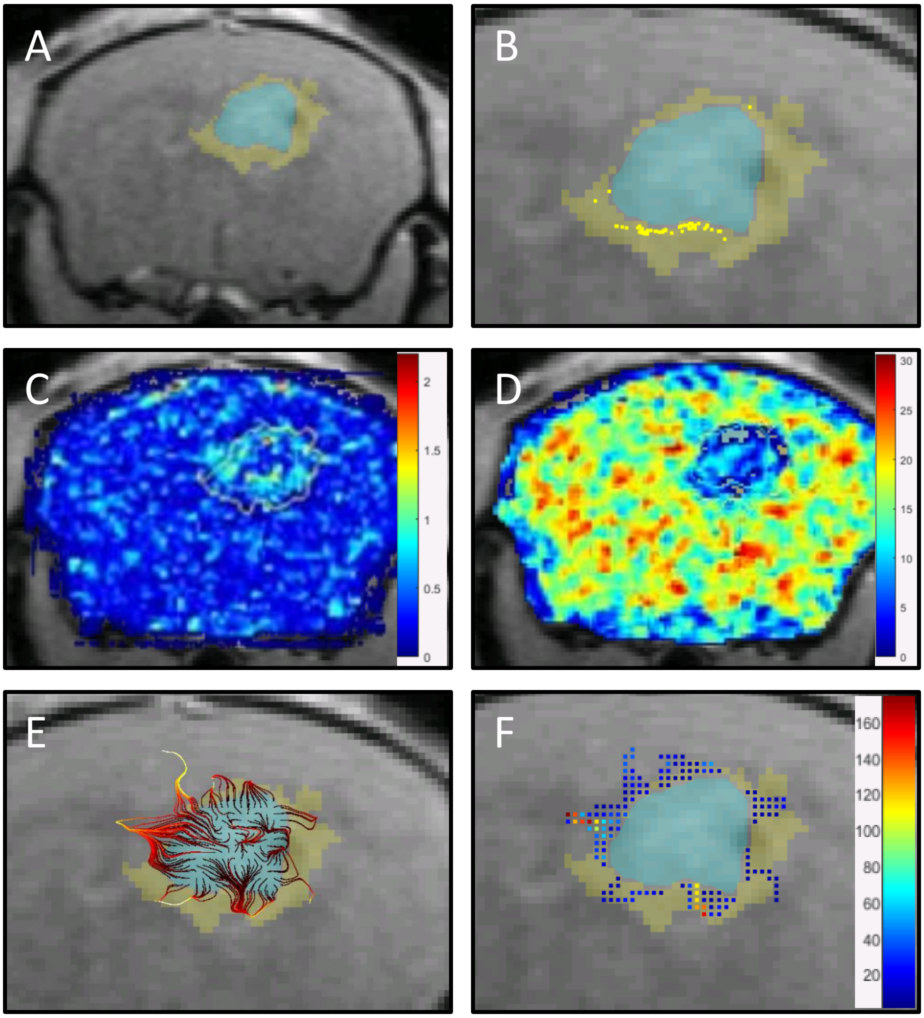


**Supplemental Dataset 1 – GL261 tumors - Mouse 5 – Location 1.** (A) Tumor (light blue highlight) and contrast-enhanced parenchyma (light yellow highlight) converted to masked regions overlaid a T1-weighted MRI (Scale bar =1mm) (B) Tumor (light blue highlight) and contrast-enhanced parenchyma (light yellow highlight) converted to masked regions overlaid a T1-weighted MRI with invading tumor cells (dark yellow points) (Scale bar = 500µm). C) Heat map of velocity magnitude on the whole brain with tumor (light blue) and contrast-enhanced parenchyma(yellow) outlined (Scale bar =1mm). D) Heat map of diffusion coefficient with the tumor (light blue) and contrast-enhanced parenchyma (yellow) outlined (Scale bar =1mm). E) Tumor-originating pathlines (dark red is initial location moving to ending location as bright yellow) overlaid on T1-weighted contrast-enhanced MRI with tumor (light blue highlight) and contrast-enhanced parenchyma (light yellow highlight) (Scale bar =500µm). F) Tumor-originating pathline density for each pixel with tumor (light blue highlight) and contrast-enhanced parenchyma (light yellow highlight). Dark blue and dark red represent a low and high tumor-originating pathline density, respectively (Scale bar =500µm).

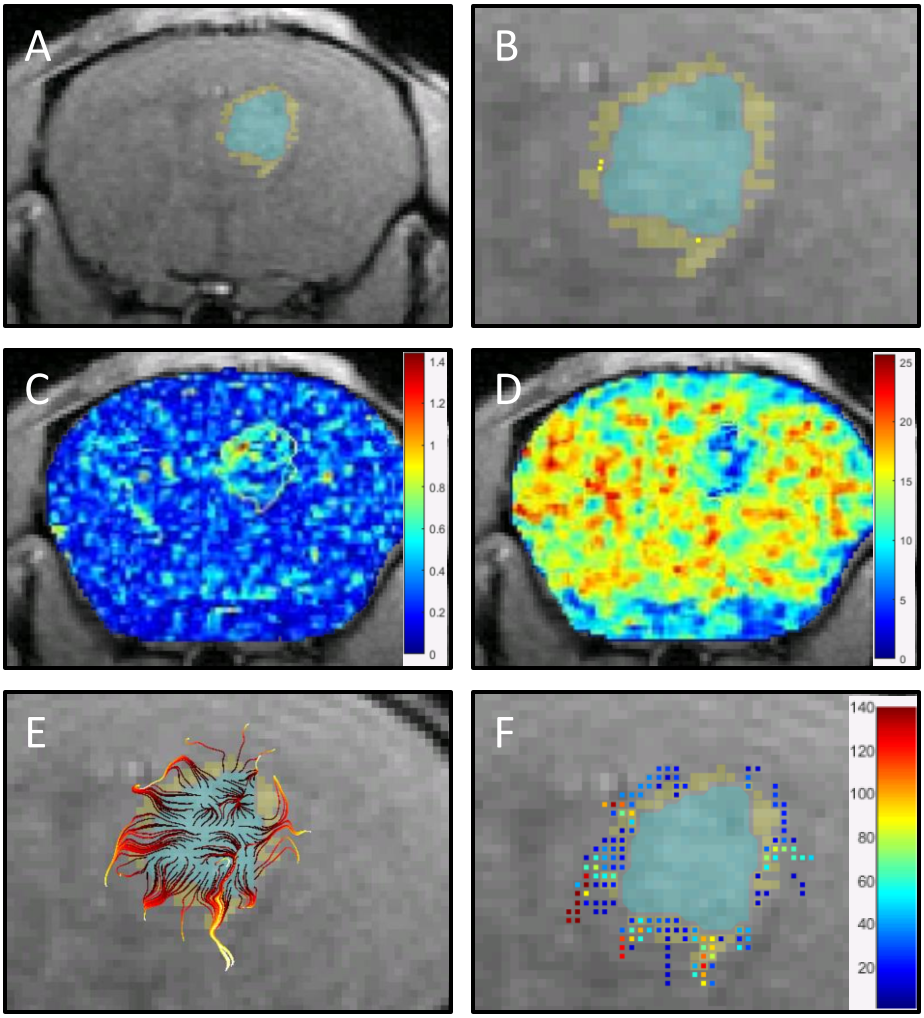


**Supplemental Dataset 1 – GL261 tumors - Mouse 5 – Location 2.** (A) Tumor (light blue highlight) and contrast-enhanced parenchyma (light yellow highlight) converted to masked regions overlaid a T1-weighted MRI (Scale bar =1mm) (B) Tumor (light blue highlight) and contrast-enhanced parenchyma (light yellow highlight) converted to masked regions overlaid a T1-weighted MRI with invading tumor cells (dark yellow points) (Scale bar = 500µm). C) Heat map of velocity magnitude on the whole brain with tumor (light blue) and contrast-enhanced parenchyma(yellow) outlined (Scale bar =1mm). D) Heat map of diffusion coefficient with the tumor (light blue) and contrast-enhanced parenchyma (yellow) outlined (Scale bar =1mm). E) Tumor-originating pathlines (dark red is initial location moving to ending location as bright yellow) overlaid on T1-weighted contrast-enhanced MRI with tumor (light blue highlight) and contrast-enhanced parenchyma (light yellow highlight) (Scale bar =500µm). F) Tumor-originating pathline density for each pixel with tumor (light blue highlight) and contrast-enhanced parenchyma (light yellow highlight). Dark blue and dark red represent a low and high tumor-originating pathline density, respectively (Scale bar =500µm).

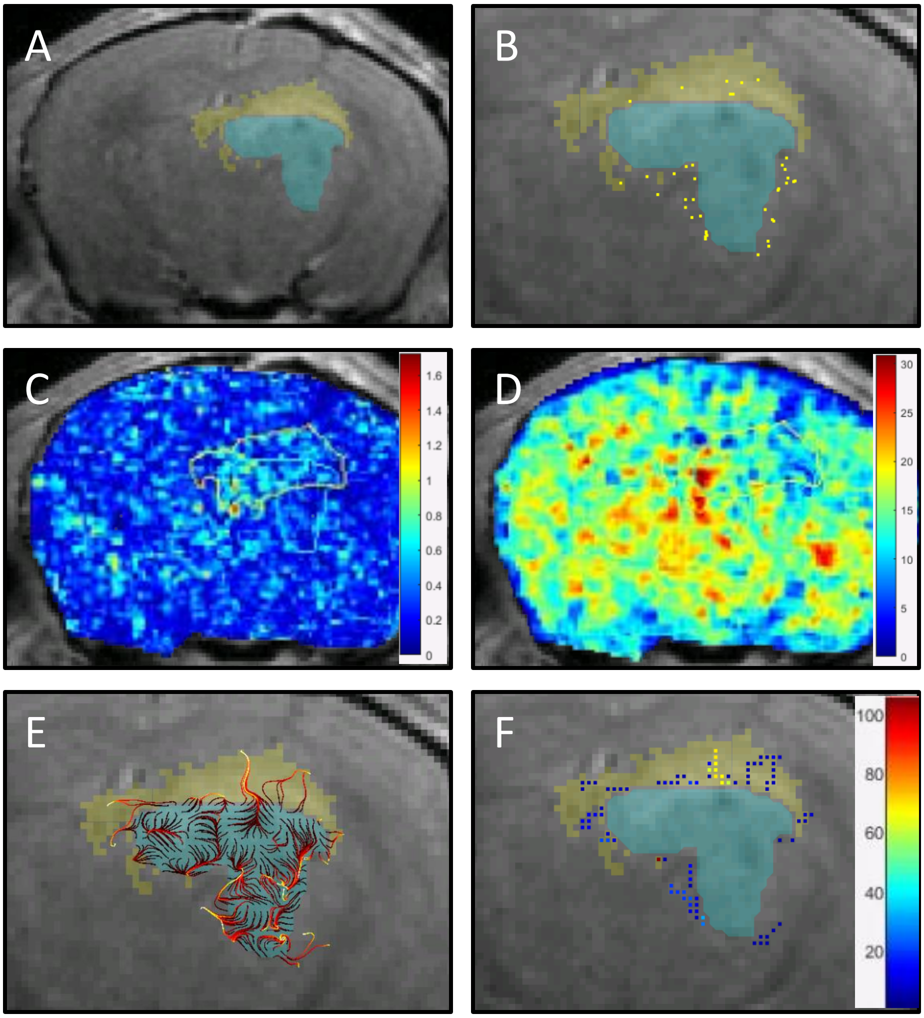


**Supplemental Dataset 1 – GL261 tumors - Mouse 7 – Location 1.** (A) Tumor (light blue highlight) and contrast-enhanced parenchyma (light yellow highlight) converted to masked regions overlaid a T1-weighted MRI (Scale bar =1mm) (B) Tumor (light blue highlight) and contrast-enhanced parenchyma (light yellow highlight) converted to masked regions overlaid a T1-weighted MRI with invading tumor cells (dark yellow points) (Scale bar = 500µm). C) Heat map of velocity magnitude on the whole brain with tumor (light blue) and contrast-enhanced parenchyma(yellow) outlined (Scale bar =1mm). D) Heat map of diffusion coefficient with the tumor (light blue) and contrast-enhanced parenchyma (yellow) outlined (Scale bar =1mm). E) Tumor-originating pathlines (dark red is initial location moving to ending location as bright yellow) overlaid on T1-weighted contrast-enhanced MRI with tumor (light blue highlight) and contrast-enhanced parenchyma (light yellow highlight) (Scale bar =500µm). F) Tumor-originating pathline density for each pixel with tumor (light blue highlight) and contrast-enhanced parenchyma (light yellow highlight). Dark blue and dark red represent a low and high tumor-originating pathline density, respectively (Scale bar =500µm).

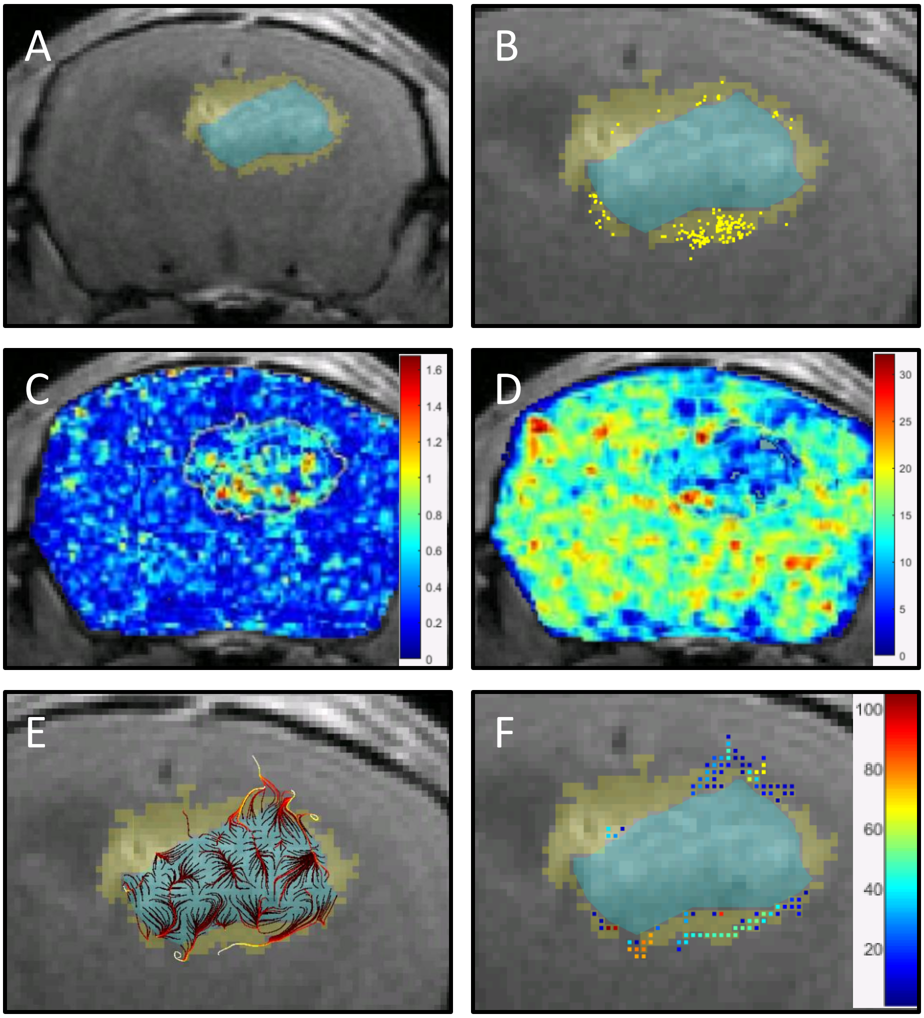


**Supplemental Dataset 1 – GL261 tumors - Mouse 7 – Location 2.** (A) Tumor (light blue highlight) and contrast-enhanced parenchyma (light yellow highlight) converted to masked regions overlaid a T1-weighted MRI (Scale bar =1mm) (B) Tumor (light blue highlight) and contrast-enhanced parenchyma (light yellow highlight) converted to masked regions overlaid a T1-weighted MRI with invading tumor cells (dark yellow points) (Scale bar = 500μm). C) Heat map of velocity magnitude on the whole brain with tumor (light blue) and contrast-enhanced parenchyma(yellow) outlined (Scale bar =1mm). D) Heat map of diffusion coefficient with the tumor (light blue) and contrast-enhanced parenchyma (yellow) outlined (Scale bar =1mm). E) Tumor-originating pathlines (dark red is initial location moving to ending location as bright yellow) overlaid on T1-weighted contrast-enhanced MRI with tumor (light blue highlight) and contrast-enhanced parenchyma (light yellow highlight) (Scale bar =500μm). F) Tumor-originating pathline density for each pixel with tumor (light blue highlight) and contrast-enhanced parenchyma (light yellow highlight). Dark blue and dark red represent a low and high tumor-originating pathline density, respectively (Scale bar =500μm).

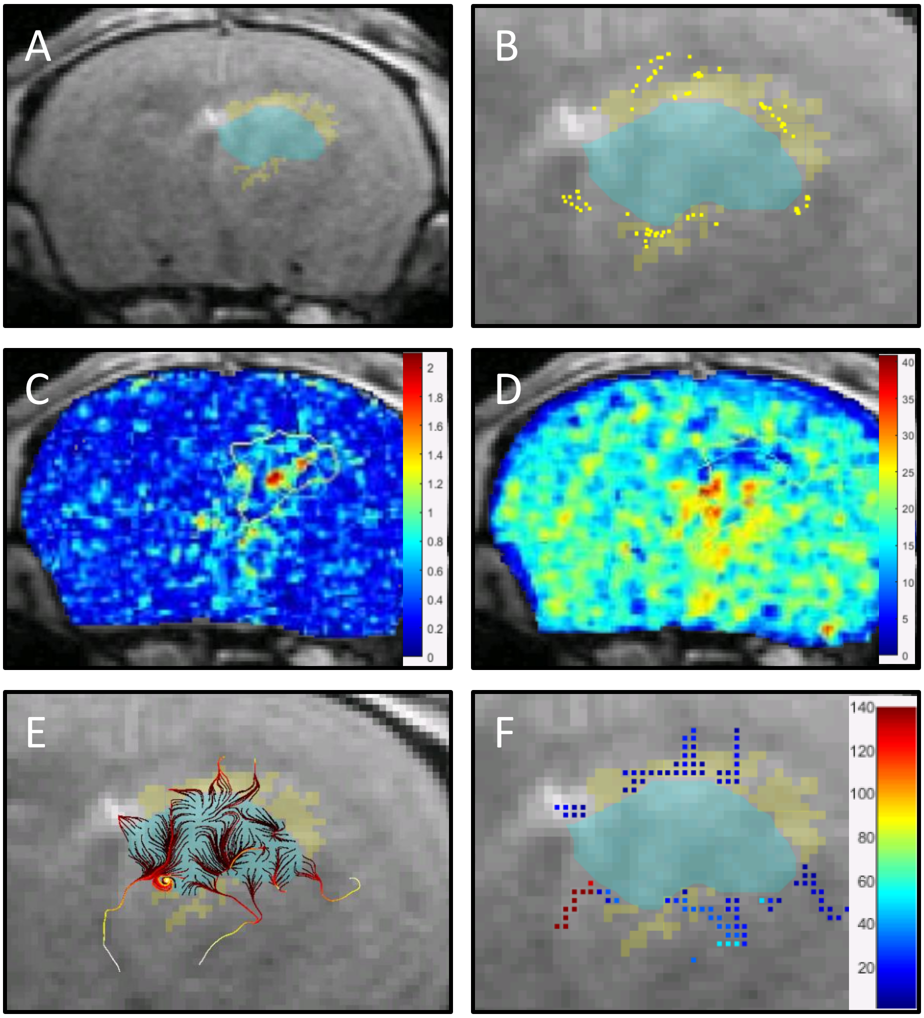


**Supplemental Dataset 1 – GL261 tumors - Mouse 10 – Location 1.** (A) Tumor (light blue highlight) and contrast-enhanced parenchyma (light yellow highlight) converted to masked regions overlaid a T1-weighted MRI (Scale bar =1mm) (B) Tumor (light blue highlight) and contrast-enhanced parenchyma (light yellow highlight) converted to masked regions overlaid a T1-weighted MRI with invading tumor cells (dark yellow points) (Scale bar = 500µm). C) Heat map of velocity magnitude on the whole brain with tumor (light blue) and contrast-enhanced parenchyma(yellow) outlined (Scale bar =1mm). D) Heat map of diffusion coefficient with the tumor (light blue) and contrast-enhanced parenchyma (yellow) outlined (Scale bar =1mm). E) Tumor-originating pathlines (dark red is initial location moving to ending location as bright yellow) overlaid on T1-weighted contrast-enhanced MRI with tumor (light blue highlight) and contrast-enhanced parenchyma (light yellow highlight) (Scale bar =500µm). F) Tumor-originating pathline density for each pixel with tumor (light blue highlight) and contrast-enhanced parenchyma (light yellow highlight). Dark blue and dark red represent a low and high tumor-originating pathline density, respectively (Scale bar =500µm).
