## Supplemental Dataset 2 for "Interstitial fluid transport dynamics predict glioblastoma invasion and progression"

### Supplemental Dataset 2: Image analysis on implanted G34 tumors with progression

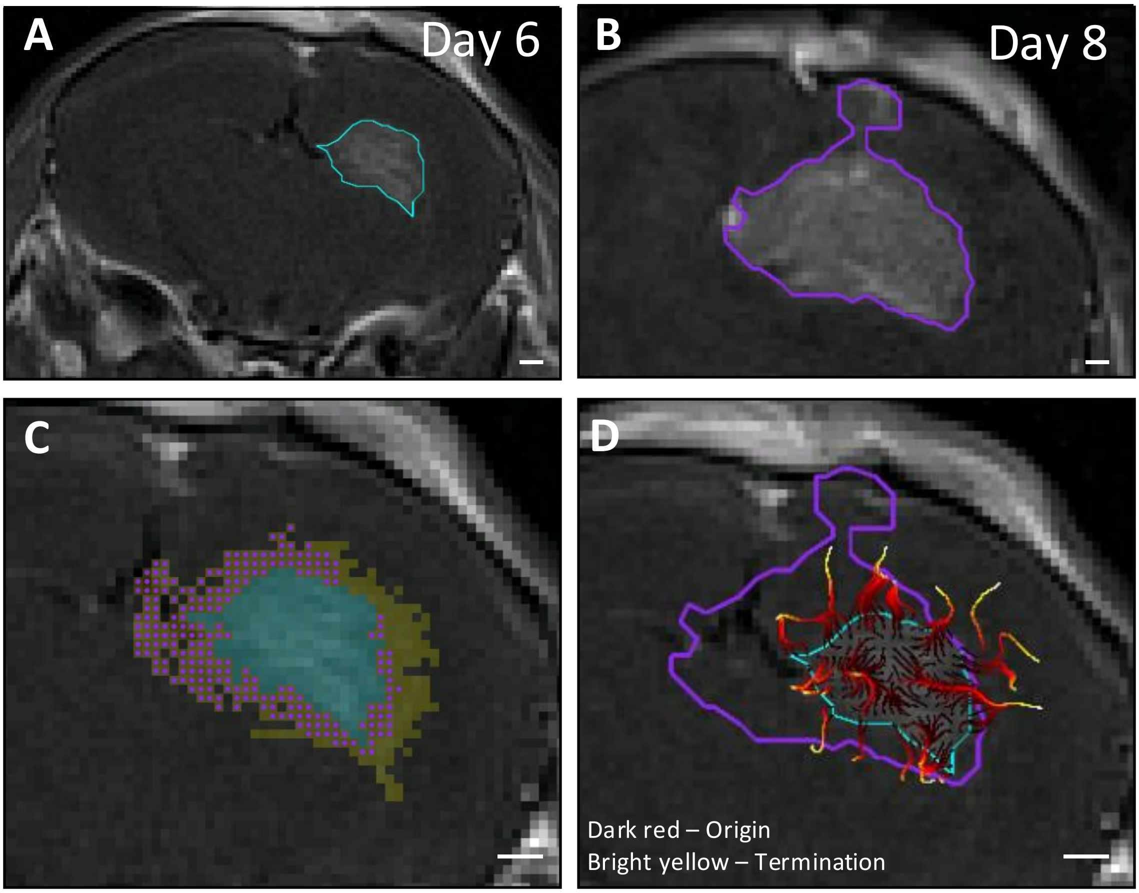


**Supplemental Dataset 2 – G34 tumors – Mouse 9.** A) Day 6 tumor boundary (cyan) as identified on T1-weighted contrast enhanced MR image. B) Day 8 tumor boundary (purple) as identified on a T1-contrast-enhanced MR image in the same tumor. C) Pixels displaying progression (purple points) within the contrast-enhancing parenchyma (yellow pixels) identified by Day 6 DCE-MRI. Pixels containing tumor on Day 8 (purple points) within the contrast-enhancing parenchyma of Day 6 (yellow pixels) are classified as "Progression," whereas all remaining contrast-enhancing pixels in the parenchyma are classified as "No Progression." D) Day 8 MRI is overlaid with Day 8 tumor boundary (magenta), Day 6 tumor boundary (cyan) and Day 6 tumor-originating pathlines (dark red to yellow transition indicates direction of pathline origination to termination) (all scale bars = 500μm).

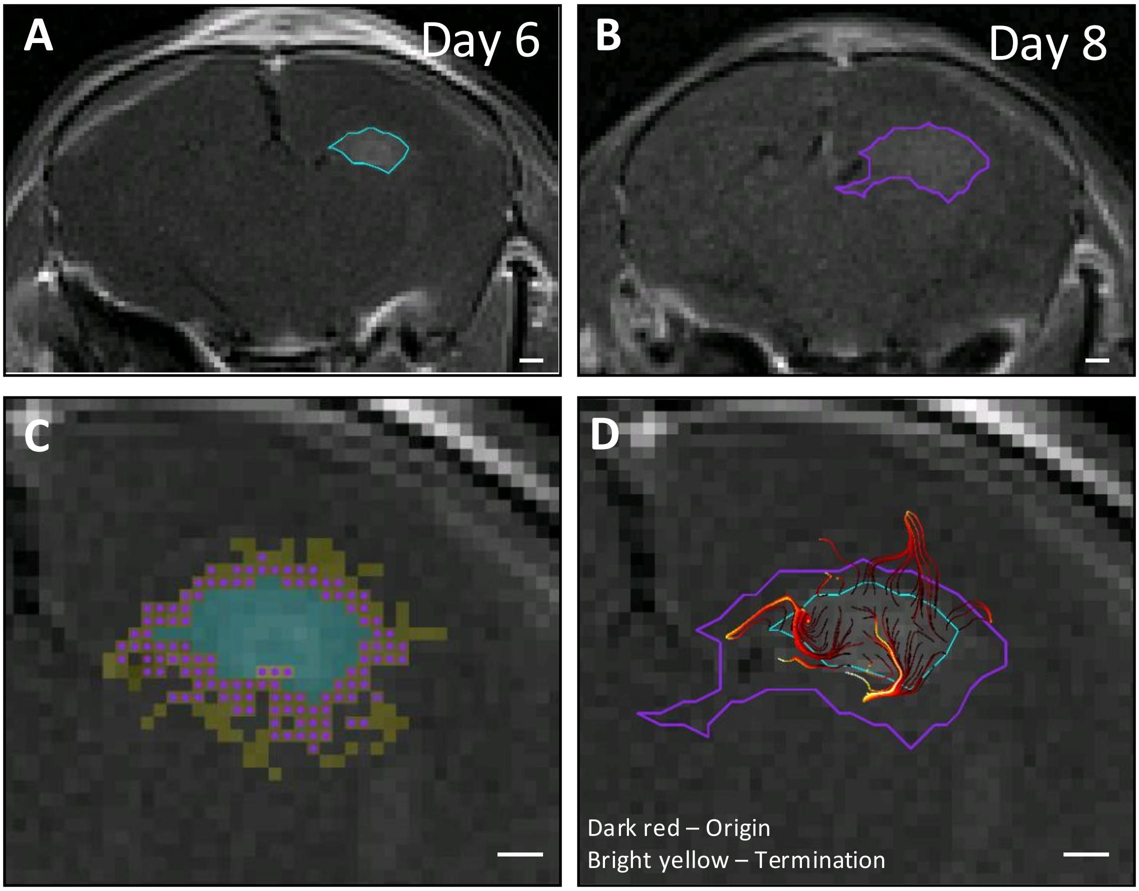


**Supplemental Dataset 2 – G34 tumors – Mouse 10.** A) Day 6 tumor boundary (cyan) as identified on T1-weighted contrast enhanced MR image. B) Day 8 tumor boundary (purple) as identified on a T1-contrast-enhanced MR image in the same tumor. C) Pixels displaying progression (purple points) within the contrast-enhancing parenchyma (yellow pixels) identified by Day 6 DCE-MRI. Pixels containing tumor on Day 8 (purple points) within the contrast-enhancing parenchyma of Day 6 (yellow pixels) are classified as "Progression," whereas all remaining contrast-enhancing pixels in the parenchyma are classified as "No Progression." D) Day 8 MRI is overlaid with Day 8 tumor boundary (magenta), Day 6 tumor boundary (cyan) and Day 6 tumor-originating pathlines (dark red to yellow transition indicates direction of pathline origination to termination) (all scale bars = 500μm).

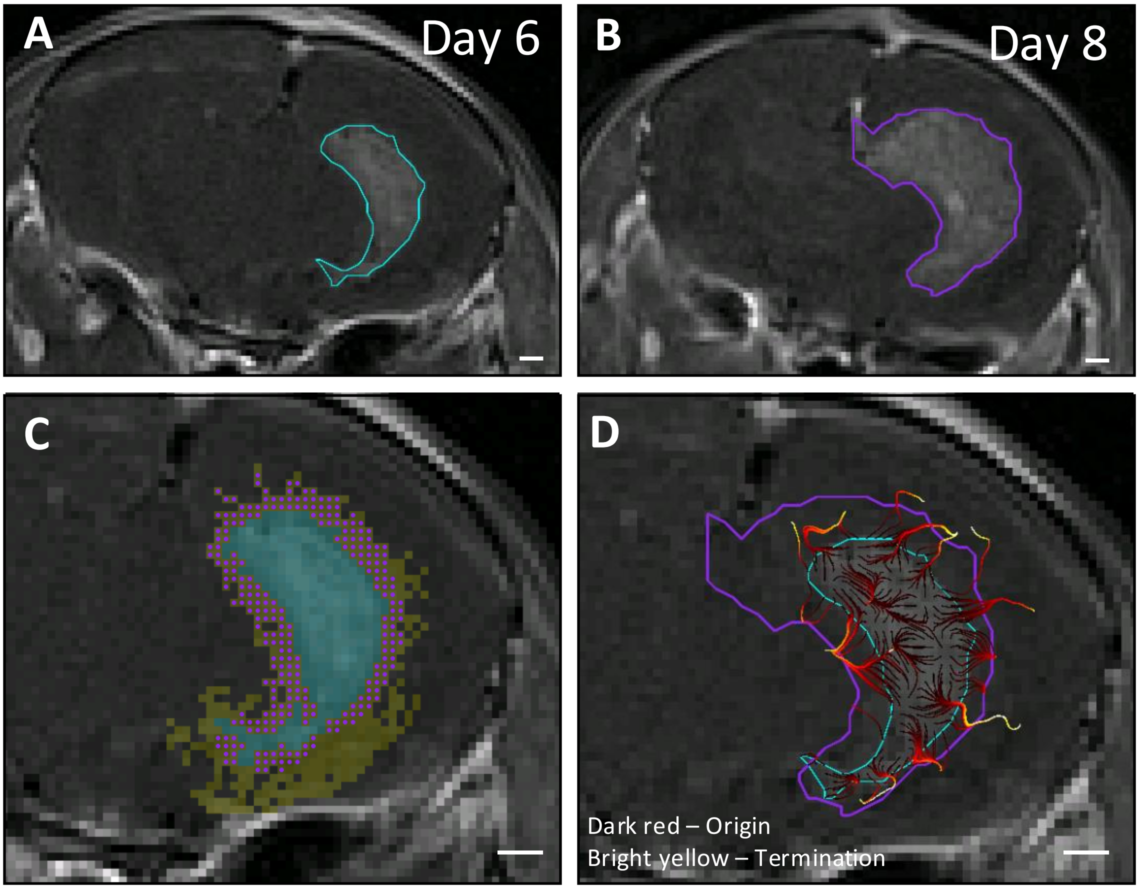


**Supplemental Dataset 2 – G34 tumors – Mouse 11.** A) Day 6 tumor boundary (cyan) as identified on T1-weighted contrast enhanced MR image. B) Day 8 tumor boundary (purple) as identified on a T1-contrast-enhanced MR image in the same tumor. C) Pixels displaying progression (purple points) within the contrast-enhancing parenchyma (yellow pixels) identified by Day 6 DCE-MRI. Pixels containing tumor on Day 8 (purple points) within the contrast-enhancing parenchyma of Day 6 (yellow pixels) are classified as "Progression," whereas all remaining contrast-enhancing pixels in the parenchyma are classified as "No Progression." D) Day 8 MRI is overlaid with Day 8 tumor boundary (magenta), Day 6 tumor boundary (cyan) and Day 6 tumor-originating pathlines (dark red to yellow transition indicates direction of pathline origination to termination) (all scale bars = 500μm).

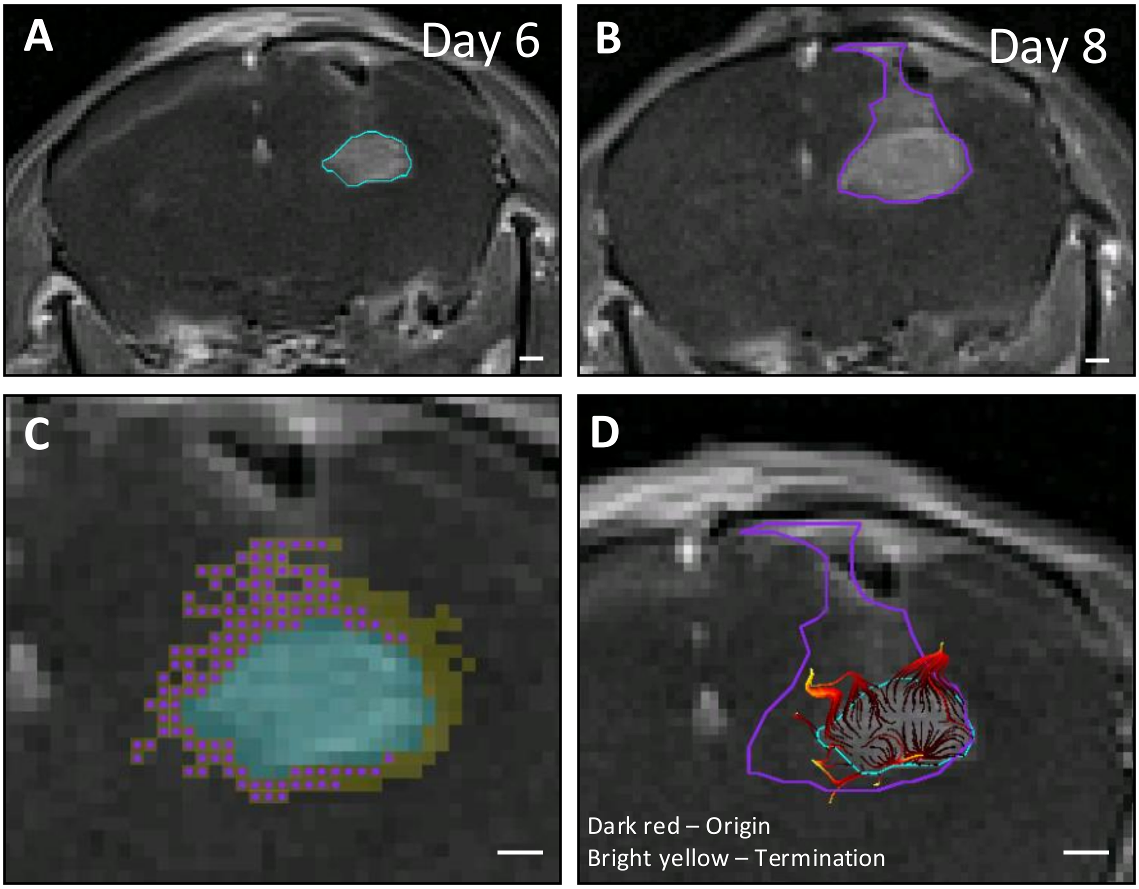


**Supplemental Dataset 2 – G34 tumors – Mouse 12.** A) Day 6 tumor boundary (cyan) as identified on T1-weighted contrast enhanced MR image. B) Day 8 tumor boundary (purple) as identified on a T1-contrast-enhanced MR image in the same tumor. C) Pixels displaying progression (purple points) within the contrast-enhancing parenchyma (yellow pixels) identified by Day 6 DCE-MRI. Pixels containing tumor on Day 8 (purple points) within the contrast-enhancing parenchyma of Day 6 (yellow pixels) are classified as "Progression," whereas all remaining contrast-enhancing pixels in the parenchyma are classified as "No Progression." D) Day 8 MRI is overlaid with Day 8 tumor boundary (magenta), Day 6 tumor boundary (cyan) and Day 6 tumor-originating pathlines (dark red to yellow transition indicates direction of pathline origination to termination) (all scale bars = 500µm).

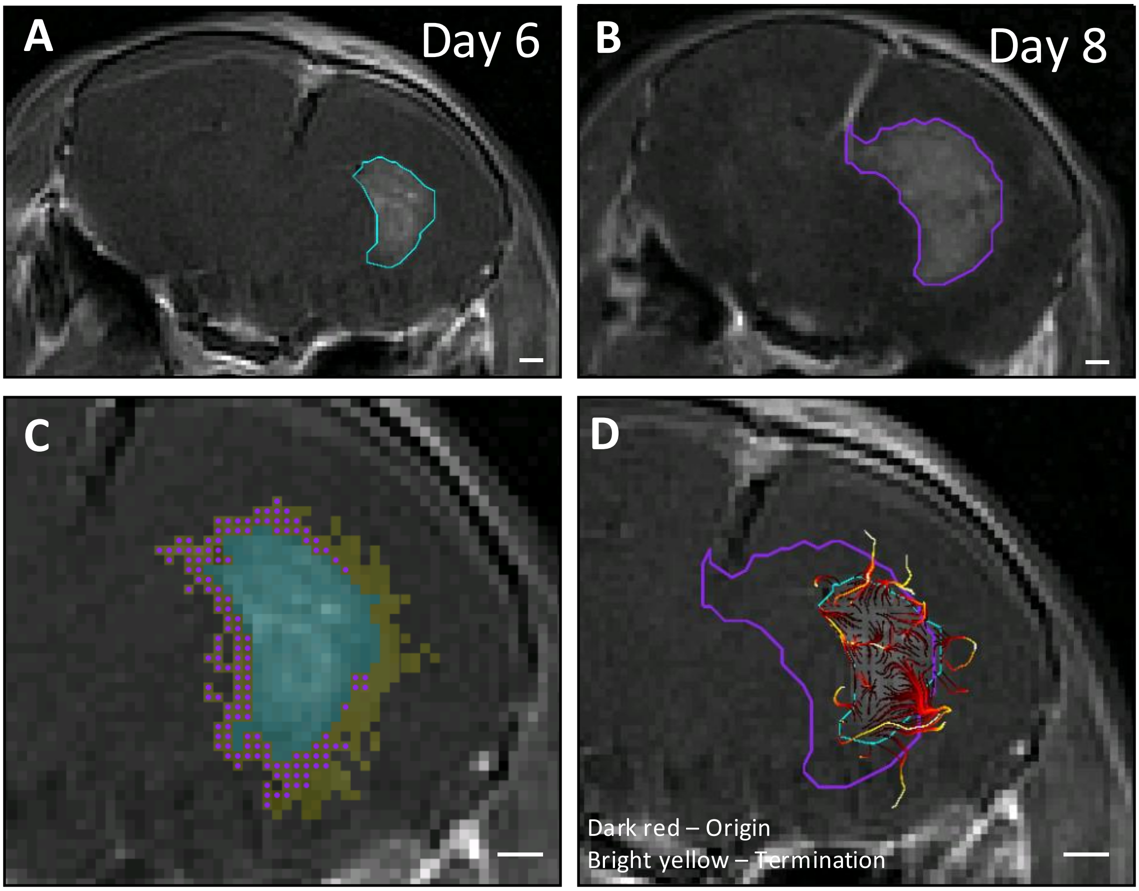


**Supplemental Dataset 2 – G34 tumors – Mouse 13.** A) Day 6 tumor boundary (cyan) as identified on T1-weighted contrast enhanced MR image. B) Day 8 tumor boundary (purple) as identified on a T1-contrast-enhanced MR image in the same tumor. C) Pixels displaying progression (purple points) within the contrast-enhancing parenchyma (yellow pixels) identified by Day 6 DCE-MRI. Pixels containing tumor on Day 8 (purple points) within the contrast-enhancing parenchyma of Day 6 (yellow pixels) are classified as "Progression," whereas all remaining contrast-enhancing pixels in the parenchyma are classified as "No Progression." D) Day 8 MRI is overlaid with Day 8 tumor boundary (magenta), Day 6 tumor boundary (cyan) and Day 6 tumor-originating pathlines (dark red to yellow transition indicates direction of pathline origination to termination) (all scale bars = 500μm).

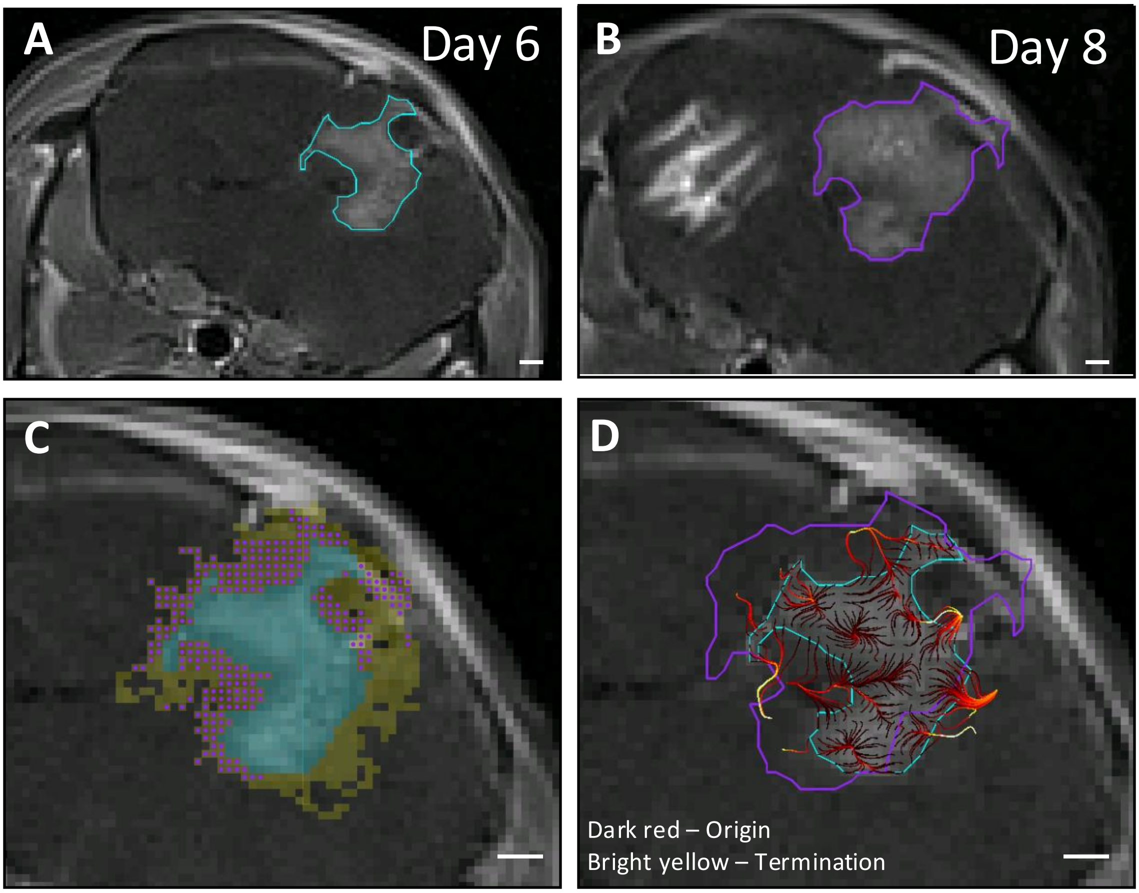


**Supplemental Dataset 2 – G34 tumors – Mouse 14.** A) Day 6 tumor boundary (cyan) as identified on T1-weighted contrast enhanced MR image. B) Day 8 tumor boundary (purple) as identified on a T1-contrast-enhanced MR image in the same tumor. C) Pixels displaying progression (purple points) within the contrast-enhancing parenchyma (yellow pixels) identified by Day 6 DCE-MRI. Pixels containing tumor on Day 8 (purple points) within the contrast-enhancing parenchyma of Day 6 (yellow pixels) are classified as "Progression," whereas all remaining contrast-enhancing pixels in the parenchyma are classified as "No Progression." D) Day 8 MRI is overlaid with Day 8 tumor boundary (magenta), Day 6 tumor boundary (cyan) and Day 6 tumor-originating pathlines (dark red to yellow transition indicates direction of pathline origination to temination) (all scale bars = 500μm).
